# Novel Pure αVβ3 Integrin Antagonists That Do Not Induce Receptor Extension, Prime the Receptor, or Enhance Angiogenesis at Low Concentrations

**DOI:** 10.1101/620765

**Authors:** Jihong Li, Yoshiyuki Fukase, Yi Shang, Wei Zou, José M. Muñoz-Félix, Lorena Buitrago, Lorena Buitrago, Johannes van Agthoven, Yixiao Zhang, Ryoma Hara, Yuta Tanaka, Rei Okamoto, Takeshi Yasui, Takashi Nakahata, Toshihiro Imaeda, Kazuyoshi Aso, Yuchen Zhou, Charles Locuson, Dragana Nesic, Mark Duggan, Junichi Takagi, Roger D. Vaughan, Thomas Walz, Kairbaan Hodivala-Dilke, Steven L. Teitelbaum, M. Amin Arnaout, Marta Filizola, Michael A. Foley, Barry S. Coller

## Abstract

The integrin αVβ3 receptor has been implicated in several important diseases, but no αVβ3 antagonists are approved for human therapy. One possible limitation of current small-molecule antagonists is their ability to induce a major conformational change in the receptor that induces it to adopt a high-affinity ligand-binding state. In response, we used structural inferences from a pure peptide antagonist to design the small-molecule pure antagonists TDI-4161 and TDI-3761. Both compounds inhibit αVβ3-mediated cell adhesion to αVβ3 ligands, but do not induce the conformational change as judged by antibody binding, electron microscopy, X-ray crystallography, and receptor priming studies. Both compounds demonstrated the favorable property of inhibiting bone resorption *in vitro*, supporting potential value in treating osteoporosis. Neither, however, had the unfavorable property of the αVβ3 antagonist cilengitide of paradoxically enhancing aortic sprout angiogenesis at concentrations below its IC_50_, which correlates with cilengitide’s enhancement of tumor growth *in vivo*.

**Significance Statement:** αVβ3 is a potential therapeutic target for several important human diseases, but there are currently no αVβ3 antagonists approved for human therapy. Current candidates are primarily based on the Arg-Gly-Asp (RGD) motif and act as partial agonists in that they induce αVβ3 to undergo a conformational change that converts it into a high-affinity ligand-binding state. We have used structure-guided design to produce pure small-molecule αVβ3 antagonists that do not induce the conformational change as judged by protein crystallography, electron microscopy, and receptor priming. These compounds inhibit αVβ3-mediated bone resorption *in vitro*, but unlike the partial agonist cilengitide, do not enhance angiogenesis at low doses, a property that correlates with low-dose cilengitide’s enhancement of tumor growth *in vivo*. These pure αVβ3 antagonists can help define αVβ3’s role in animal models. If they demonstrate benefits over partial agonists in these model systems, they may be appropriate to consider for human therapy.

## Introduction

The β3 integrin family, composed of αVβ3 and αIIbβ3, has been intensively studied and many of the details about integrin structure and function in general have come from studies of this family. αIIbβ3 is specific for megakaryocytes and platelets, plays a key role in hemostasis and thrombosis,^1^ and is a validated drug target to prevent thrombosis.^2,3^ αVβ3 is expressed on osteoclasts^4^ and variably expressed on other tissues in response to different stimuli, including hypoxia and other cellular stresses, most notably on neovascular endothelial cells.^5,6^ It plays an important role in bone resorption and has been implicated in contributing to a broad range of pathological processes. These include osteoporosis,^7,8^ sickle cell disease vaso-occlusion,^9–11^ tumor angiogenesis,^6,12^ metastasis,^5^ tumor-induced bone resorption,^13^ herpes simplex and hantavirus viral invasion,^14–16^ disruption of glomerular barrier function,^17,18^ dermal and hepatic fibrosis,^19–24^ acute myelogenous leukemia,^25^ post-cardiac transplant coronary vasculopathy,^26,27^ bone resorption by multiple myeloma plasma cells,^28^ supravalvular aortic stenosis associated with Williams syndrome,^29^ Crohn’s disease strictures,^30^ and T-cell lymphoma.^31,32^ Despite the potential clinical utility of inhibiting αVβ3, there are no approved drugs or biologics targeting this receptor.^33^ Human studies with the RGD-containing cyclic pentapeptide Arg-Gly-Asp-{D-Phe}-{N-*methy*l-Val}, cilengitide, which inhibits αVβ3 and αVβ5, failed to demonstrate efficacy for treating glioblastoma,^34^ although it may be beneficial with tumors expressing high levels of αVβ3.^35^ In contrast, the RGD-based αVβ3 antagonist MK-429 showed promising biomarker improvements and an increase in bone mineral density, as well as a favorable safety profile, when administered to postmenopausal women for 12 months to prevent the progression of osteoporosis.^36^ It also showed favorable effects on biomarkers of bone turnover in 21 men with bone metastases from prostate carcinoma,^37^ but its clinical development was stopped for unknown reasons.

The binding of ligands or RGD-based integrin antagonists to the closely related αIIbβ3 receptor initiates major conformational changes in the receptor.^38–43^ The interaction of the ligand carboxyl with the metal ion in the Metal Ion-Dependent Adhesion Site (MIDAS) in β3 triggers the conformational change by inducing the neighboring β1-α1 loop to move toward the MIDAS. This relatively subtle movement leads to a major swing-out motion of the hybrid domain that exposes new epitopes for monoclonal antibodies (mAbs) and induces the receptor to extend and adopt a high-affinity ligand-binding state.^39–44^ Thus, the RGD-based αIIbβ3 antagonists are partial agonists and can under certain experimental conditions actually prime the receptor to bind ligand in the absence of an activating agent.^42,44–48^ This effect has been hypothesized to explain the increased mortality in patients treated with the oral RGD-based αIIbβ3 antagonists that did not gain approval for human use.^49–51^ The conformational changes in αIIbβ3 induced by the approved RGD-based partial agonist drugs eptifibatide and tirofiban are also hypothesized to account for the development of thrombocytopenia in ~0.5-1% of treated patients by exposing αIIbβ3 epitopes to which some individuals have pre-formed antibodies, resulting in antibody coating of platelets and their rapid removal from the circulation.^49,52,53^

The binding of RGD-based antagonists to αVβ3 produces similar changes in metal-ion coordination to those observed with the RGD-based antagonists to αIIbβ3^54^ and also exposes a Ligand-Induced Binding Site (LIBS) on the β3 PSI domain for mAb AP5.^55–57^ This partial agonist activity may contribute to inducing receptor extension,^58^ priming the receptor to bind ligand at low doses *in vitro*,^59^ enhancing angiogenesis *ex vivo* at sub-IC_50_ concentrations,^57^ and activating osteoclast signaling like that produced by natural ligands.^60^ Many of the animal studies supporting a role for αVβ3 in the pathophysiology of disease have included evidence that cilengitide improves outcome. In the case of hepatic fibrosis, however, cilengitide paradoxically produced increased collagen deposition due to activation of hepatic stellate cells despite its positive impact on cell-based assays.^61^ The enhancement of *ex vivo* angiogenesis by cilengitide at sub-IC_50_ concentrations is of particular concern because it correlated with paradoxical enhancement of *in vivo* tumor formation in mice at sub-IC_50_ concentrations and thus, although speculative, may contribute to the lack of clinical efficacy of cilengitide in treating glioblastoma.^57^ Thus, it is important to assess whether a pure αVβ3 antagonist, that is, one that blocks the receptor without inducing the conformational change, would have therapeutic benefits that have not been observed with the current partial agonist small molecules.

Arnaout’s group reported that whereas the RGD-containing native fibronectin fragment FN10 is a partial agonist of αVβ3, a mutant form of the peptide (hFN10) acts as a pure antagonist. This change was ascribed to the substitution of a Trp residue for a Ser immediately after the RGD sequence because the Trp forms a *π*-*π* interaction with β3 Tyr122 on the β1-α1 loop, thus preventing the latter’s movement toward the MIDAS, a key element in triggering the conformational change.^62^ The importance of interacting with Tyr122 to prevent the conformational change in αVβ3 is also supported by studies demonstrating that non-enzymatic conversion of Asn to isoAsp in the GNGRG sequence in fibronectin repeat 5 results in the repeat developing high affinity for αVβ3; the cyclic peptide CisoDGRC is reported to retain this high affinity without apparently inducing the conformational change in αVβ3^63,64^ because the C1 of the peptide interacts via its N-terminus with the Tyr122 carbonyl in β3.^63,64^

Based on Arnaout’s observations, we synthetically modified the high-affinity RGD-based αVβ3 antagonist MK-429 so as to establish a *π*-*π* interaction with β3 Tyr122, guided by a three-dimensional molecular model of MK-429’s interaction with αVβ3 refined by molecular dynamics (MD) simulations. We searched for compounds that inhibit the adhesion of HEK-293 cells expressing αVβ3 to one of its ligands (fibrinogen), but do not trigger the activating conformational change in the receptor. We monitored the induction of the swing-out conformation in the β3 subunit by assessing the exposure of the epitope on β3 for the LIBS mAb AP5 and then confirmed the results by both protein crystallography and by assessing receptor extension and swing-out by electron microscopy. Our goal is to obtain compounds that retain the ability to inhibit αVβ3 ligand binding where it contributes to disease, as in osteoclast resorption of bone, while eliminating their ability to induce the conformational change that may prime or signal through the receptor, and thus may be responsible for enhanced angiogenesis at sub-IC_50_ concentrations. Therefore, we compared the biologic effects of our compounds that met the above criteria with those of current RGD-based compounds.

## Results

### Properties of current RGD-based αVβ3 antagonists

The RGD-based compounds cilengitide and a racemic mixture of MK-429 (i.e., containing both MK-429 enantiomers),^36,37,65–67^ were characterized by their ability to inhibit the adhesion of HEK-293 cells expressing human αVβ3 to immobilized fibrinogen (IC_50_) and their ability to induce the exposure of the epitope for mAb AP5 (EC_50_); thus, higher values for the ratio of EC_50_ to IC_50_ indicate that the compound is less able to induce the conformational change. The RGD-based compounds cilengitide and the racemate of MK-429 had IC_50_s of 29 and 3 nM, EC_50_s of 48 and 12 nM, and EC_50_/IC_50_ ratios of 1.7 and 4.0, respectively (Table 1).

**Table 1.**

|  | <u>Cilengitide</u> |  | <u>MK-429</u><br><u>racemate</u> |  | <u>TDI-3761</u> |  | <u>TDI-4161</u> |
| --- | --- | --- | --- | --- | --- | --- | --- |
| IC<sub>50</sub> (μM) for inhibition of human αVβ3-mediated cell adhesion to fibrinogen | 0.029 ± 0.040<br>(n=6) |  | 0.003 ± 0.002<br>(n=6) |  | 0.172 ± 0.066<br>(n=5) |  | 0.025 ± 0.010<br>(n=6) |
| EC<sub>50</sub> (μM) for mAb AP5 epitope exposure | 0.048 ± 0.013<br>(n=3) |  | 0.012 ± 0.001<br>(n=3) |  | >10<br>(n=3) |  | >10<br>(n=3) |
| EC<sub>50</sub>/IC<sub>50</sub> | 1.7 |  | 4.0 |  | >58 |  | >400 |
| IC<sub>50</sub> (μM) for inhibition of murine αVβ3-mediated cell adhesion to fibrinogen | 0.026 ± 0.010<br>(n=4) |  | 0.004 ± 0.001<br>(n=4) |  | 0.041 ± 0.027<br>(n=6) |  | 0.067 ± 0.046<br>(n=6) |
| IC<sub>50</sub> (μM) for inhibition of purified αVβ3 binding to penton base | 0.007 ± 0.008<br>(n=4) |  | 0.011 ± 0.005<br>(n=4) |  | 0.049 ± 0.050<br>(n=4) |  | 0.012 ± 0.005<br>(n=4) |
| IC<sub>50</sub> (μM) for inhibition of purified αVβ5 binding to vitronectin | 0.006 ± 0.004<br>(n=5) |  | 0.035 ± 0.019<br>(n=5) |  | 0.875 ± 0.348<br>(n=5) |  | 0.689 ± 0.374<br>(n=5) |

### Structure-based design of new compounds

We docked MK-429 into the site occupied by hFN10 in the crystal structure of the αVb3-hFN10 complex (PDB ID: 4MMZ) as described in Methods (Fig.1). In the top docking pose of MK-429, whose stability was confirmed by MD simulations (Supporting Information Figs. S1-S3), the tetrahydronaphthyridin moiety interacts with Asp218 of the αV subunit and the carboxylic acid moiety binds the Mg^2+^ in the β3 subunit MIDAS. However, in sharp contrast with the receptor interactions formed by hFN10, the ligand’s aromatic ring at the carboxylate end does not make an aromatic *π*-*π* stacking interaction with β3 Tyr122, thus possibly allowing for the movement of the β3 β1-α1 loop toward the MIDAS and the exposure of the AP5 epitope. Specifically, the distance between the centroids of the two aromatic rings is 7.8 Å for MK-429 compared to 4.7 Å for hFN10. Our goal, therefore, was to design a compound that retained the nanomolar affinity of MK-429 for αVβ3 while creating an aromatic interaction with β3 Tyr122 similar to that of the Trp in hFN10, thus potentially preventing the movement of the β1-α1 loop toward the MIDAS and the resulting conformational change.

**Fig. 1.**
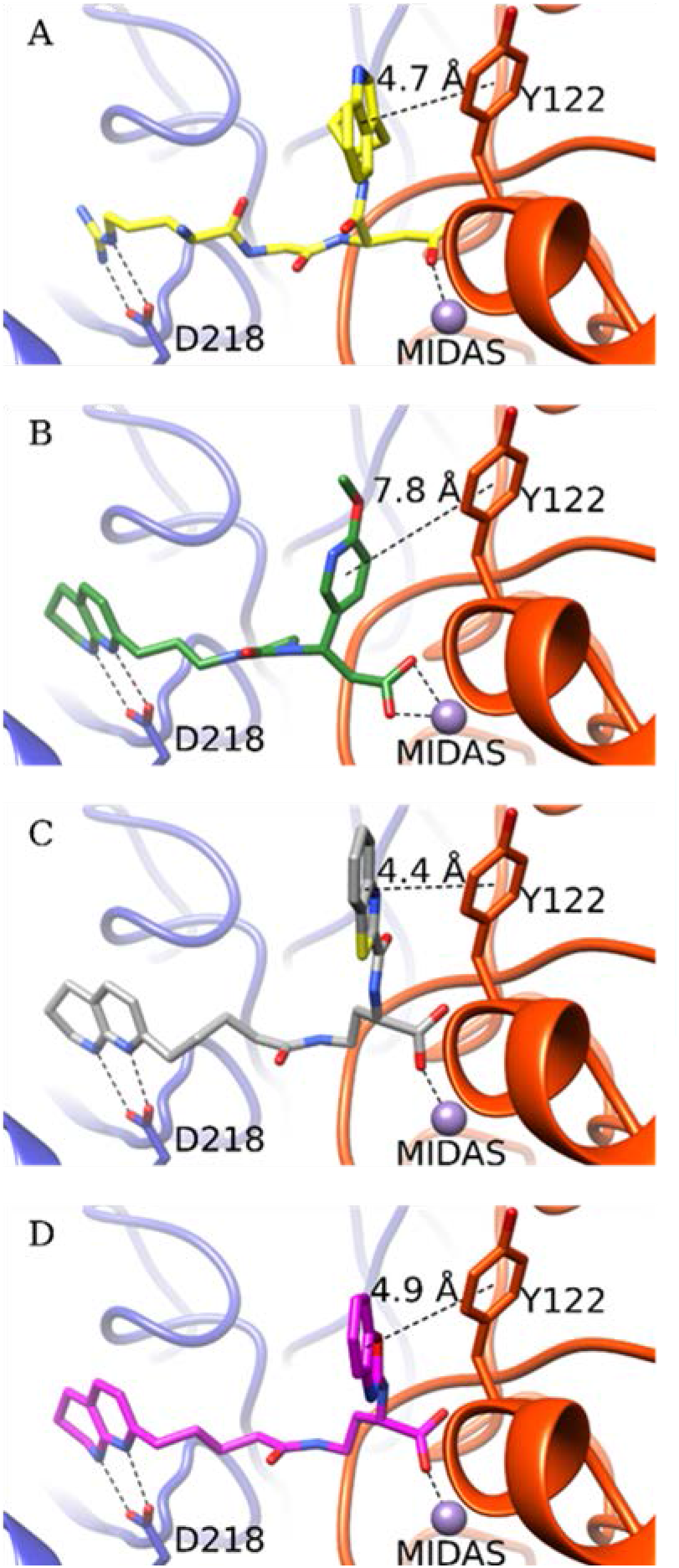
Crystal structure of the high-affinity fibronectin fragment hFN10 (A) and the predicted docking poses of MK-429 (B), TDI-4161 (C), and the *S*-enantiomer of TDI-3761 (D) in αVβ3. The αV and β3 backbones are shown in blue and red cartoon representations, respectively. Side chains of αV-Asp218 and β3-Tyr122 are shown as sticks. The MIDAS metal ion is shown as a purple sphere. The interactions between the compounds and αV-Asp218 and the MIDAS metal ion are indicated by dotted lines. Distances are reported in Å between the TyrR122 centroid *π* ring and centroids of aromatic groups at the α position of the compound’s carboxylic acid.

To design hFN10-like small molecules, we first simplified the synthesis by removing first the imidazolidinone group of TDI-806, the racemate of MK-429 (Fig. 2), yielding TDI-1366, and then the substituent at the β-position of the carboxylic acid, yielding TDI-1367, which demonstrated reduced potency, but greater selectivity in not exposing the AP5 epitope. With this simplified and selective scaffold, we undertook parallel synthesis to probe the structure-activity relationship around the amino acid moiety of TDI-1367. Lengthening the compound by one carbon yielded TDI-2668, which demonstrated reduced potency, but greater selectivity. We subsequently explored aromatic substitutions at the α and β positions of the carboxylic acid with the goal of developing *π*-*π* stacking with β3 Tyr122, and found that substitutions at both positions could increase potency, but only those at the α position preserved high selectivity, yielding the racemates TDI-3761 and TDI-3909. Both enantiomers of TDI-3761 have properties similar to those of TDI-3761 [*S*-enantiomer (TDI-4160) IC_50_ = 0.185 ± 0.042 µM (n=5); EC_50_ > 10 μM (n = 3); *R*-enantiomer (TDI-4158) IC_50_ = 0.123 ± 0.055 μM (n=5); EC_50_ > 10 μM (n = 3)], but the *S*-enantiomer of TDI-3909, TDI-4161, is more potent and equally selective (Table 1).

**Fig. 2.**
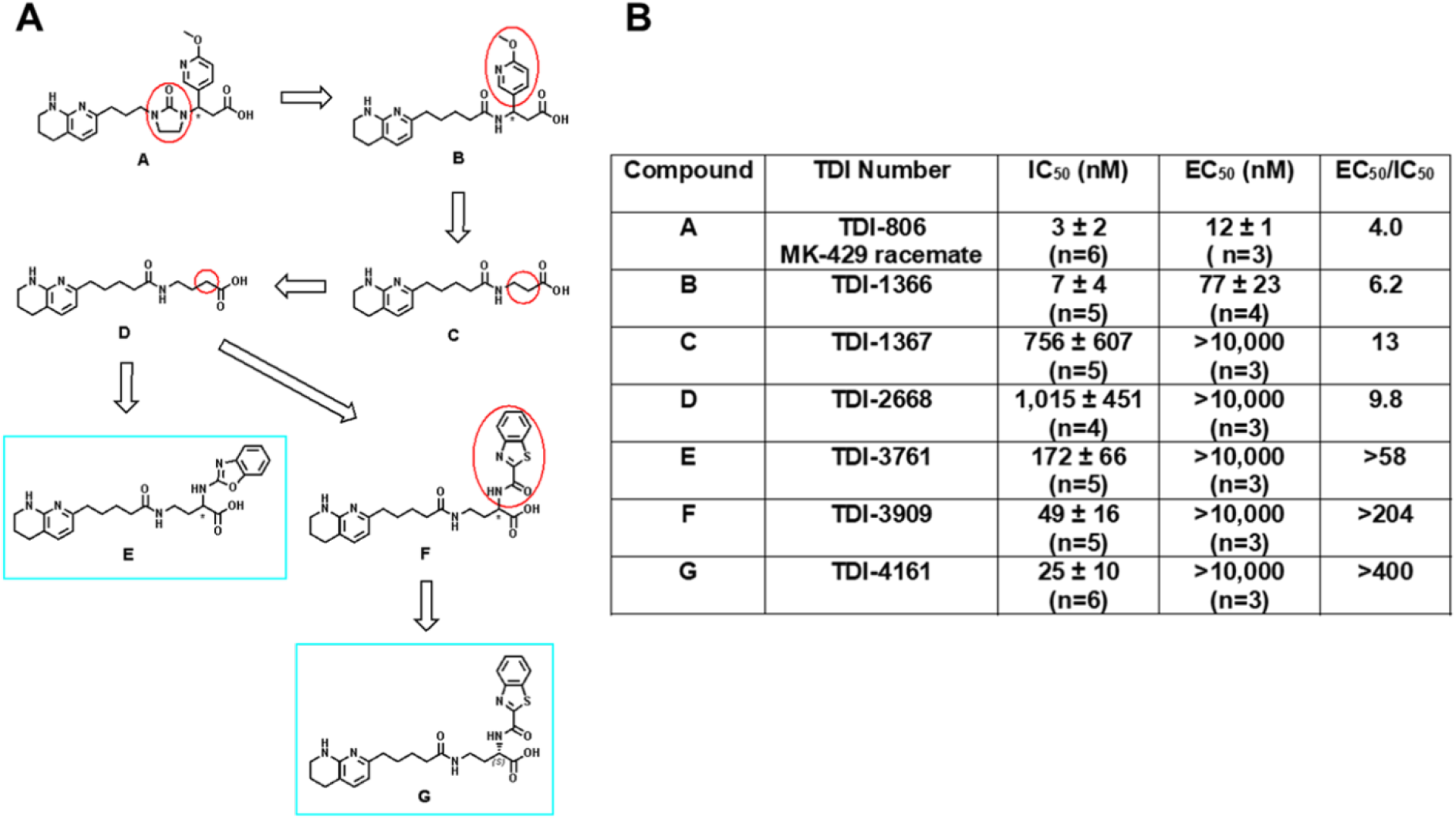
Development of TDI-4161 and TDI-3761. A. Structures. Structural modifications began with TDI-806, the racemate of MK-429 (A). The first step involved removing the imidazolinone ring, yielding TDI-1366 (B). This compound was further simplified by removing the aromatic sidechain, yielding TDI-1367(C). Increasing the length of TDI-1367 by one carbon resulted in TDI-2668 (D). The racemic compounds TDI-3761 (E) and TDI-3909 (F) were produced by adding aromatic groups in the α position of the TDI-2668 carboxylic acid. Both the *R* and *S* enantiomers of TDI-3761 had properties similar to those of TDI-3761 (see text for values), whereas TDI-4161 (G), the *S* enantiomer of TDI-3909 was more potent and equally selective when compared to TDI-4169, the *R* enantiomer (not shown). **B. Characteristics of compounds.**

TDI-3761 also inhibited the adhesion of HEK-293 cells expressing human αVβ3 to fibronectin [IC_50_ = 0.079 ± 0.010 (mean ± SD) μM; n = 3] and vitronectin (IC_50_ = 0.125 ± 0.019 μM; n = 3); comparable results for TDI-4161 were 0.042 ± 0.021 and 0.050 ± 0.017 μM (n = 3), respectively.

The dockings of TDI-4161 and the *S*-enantiomer of TDI-3761 to αVβ3 (Fig. 1; also see Methods section) show *π*-*π* interactions between the ligand’s aromatic ring at the carboxylate end and β3 Tyr122, with distances between centroids of 4.4 and 4.9 Å, respectively. The stability of this interaction in TDI-4161 binding was confirmed by MD (Fig. S4).

### Crystal structure of the αVβ3-TDI-4161 complex

To assess the validity of the docking and MD simulations of the αVβ3-TDI-4161 complex, we determined the crystal structure of the complex. A diffraction data set to 3.0 Å was obtained from αVβ3 crystals soaked with TDI-4161 in 1 mM MnCl_2_ (Table S1). Clear densities for TDI-4161 and the Mn^2+^ ions at the LIMBS, MIDAS and ADMIDAS were found at the RGD-binding pocket (Fig. 3A). As in the crystal structure of the αVβ3-hFN10 complex (Fig. 3B), the RGD-binding pocket in the αVβ3-TDI-4161 complex is lined with Tyr178 and Asp218 in αV, the MIDAS ion, and Tyr122, Arg214, and Met180 in β3. Similar to inferences from the predicted docking pose (see overlap with crystal structure in Fig. S5), the Arg-mimetic naphthyridine group of TDI-4161 hydrogen bonds αV-Asp218 and additionally makes a *π*-*π* stacking interaction with αV-Tyr178. The Asp-mimetic carboxylic group directly coordinates the metal ion at MIDAS. Significantly, the benzothiazole group forms a parallel *π*-*π* stacking interaction with β3-Tyr122, resulting in a centroid-to-centroid distance of 4.2 Å (versus 4.7 Å for the T-shaped *π*-*π* stacking between hFN10-Trp1496 and β3-Tyr122). This interaction appears to prevent the inward movement of the β1-α1 loop towards the MIDAS and consequently the large change in the F/α7 loop that leads to the activating swing-out movement of the hybrid domain.^68^ The benzothiazole group of TDI-4161 also makes an S-*π* interaction with β_3_-Met180 and a cation-*π* interaction with the guanidinium group of β_3_-Arg214. These S-*π* and cation-*π* interactions help position the benzothiazole group to face β3-Tyr122. We conclude that TDI-4161 and TDI-3761, like hFN10, most likely do not expose the AP5 epitope because the interaction with Tyr122 prevents the movement of the β1-α1 loop of the βA domain.

**Fig. 3.**
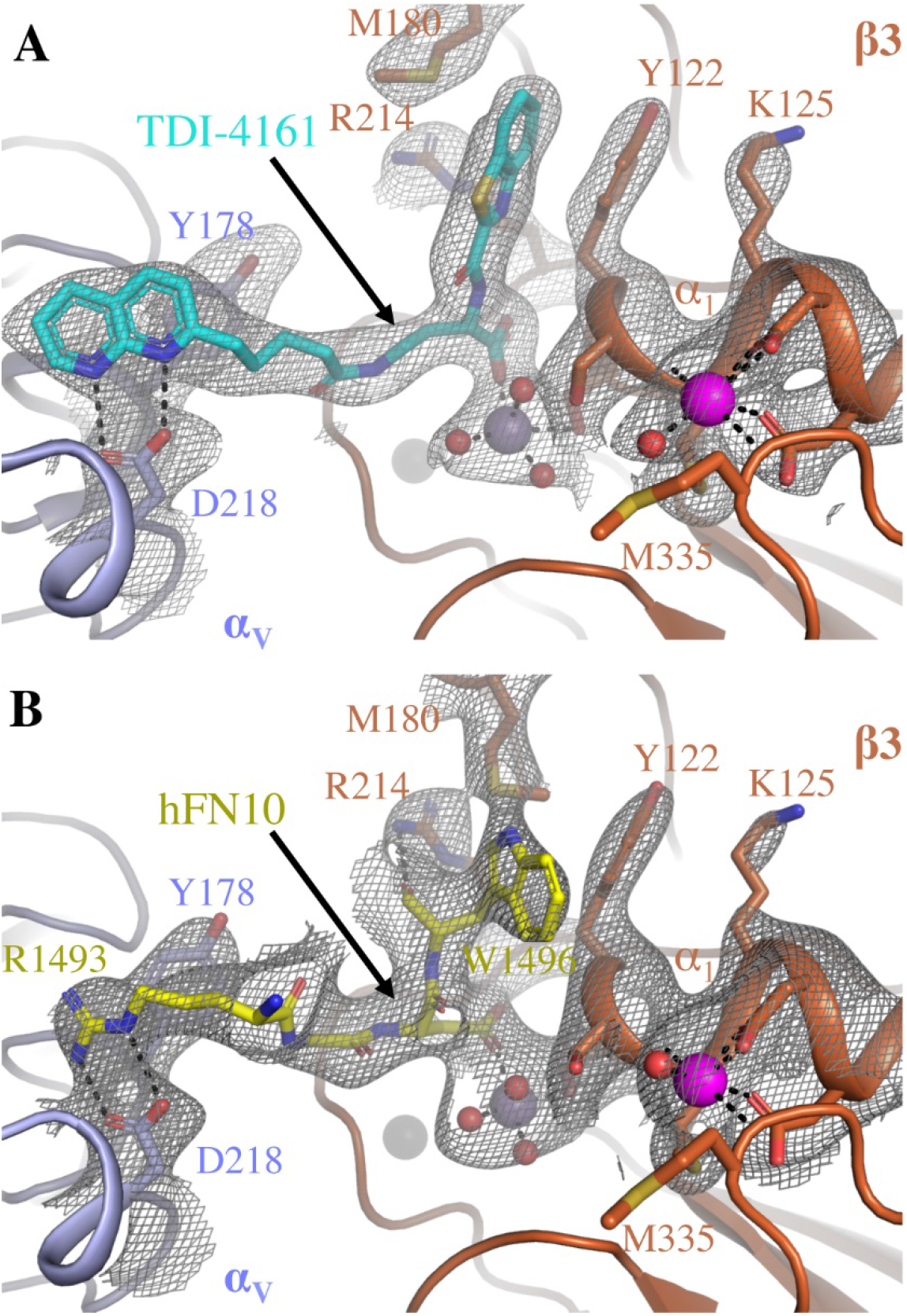
Comparison of the binding pockets of TDI-4161 in the crystal structure of the αVβ3-TDI-4161 complex and the RGDW sequence of hFN10 in the crystal structure of the αVβ3-hFN10 complex. (A) 2Fobs−Fcalc electron density map (at 1.0 σ) of TDI-4161 (shown in cyan) and TDI-4161 binding pocket of the αVβ3–TDI-4161 complex. (B) 2Fobs–Fcalc electron density map (at 1.0 σ) of the RGDW sequence of hFN10 (shown in yellow) and the RGDW binding pocket of the αVβ3–hFN10 complex. αV propeller is shown in light blue, β3A domain in copper, water molecules as red spheres and the Mn^2+^ ions at the LIMBS, MIDAS and ADMIDAS as grey, purple, and magenta spheres, respectively. TDI-4161, hFN10, and αVβ3 side-chain and backbone atoms are shown as sticks in the respective colors. Oxygen, nitrogen, and sulfur atoms are in red, blue and yellow, respectively.

Interestingly, freezing the αVβ3 ligand-binding site in the inactive conformation by bound TDI-4161 is associated with quaternary changes mainly seen in the membrane-proximal β-terminal domain (βTD). TDI-4161 binding is associated with formation of a hydrogen bond between Gln319 in the βA domain and Ser674 of the βTD, and stabilization of a glycan at Asn654 (Fig. S6), changes also seen in the αVβ3-hFN10 structure.^62^

### Electron microscopic assessment of αVβ3 conformational change

Class averages obtained with negatively stained αVβ3 in the presence of DMSO (control) or αVβ3 antagonists (all at 10 μM) are shown in Fig. S7, with the number of particles in each class indicated by the number below the average. The class averages were manually assigned to represent molecules that were in a compact-closed conformation (red border), an extended-closed conformation (blue border), or an extended-open conformation (green border). Class averages that did not show clear structural features of one of these conformations were not assigned and removed from further analysis. The total number of images analyzed ranged from 14,933-18,693, and the percentage of unassigned images ranged from ~9-20% (Figs. 4 and S7). The percentage of assigned molecules in the compact-closed conformation in the control sample (0.1% DMSO vehicle) was 77%, whereas this group constituted only 20% of the molecules in the presence of cilengitide and 27% of the molecules in the presence of MK-429. In dramatic contrast, in the presence of TDI-4161 and TDI-3761, the comparable values were 79% and 67%, respectively, which are similar to the control sample. In reciprocal fashion, just 3% of assignable molecules in the control sample were in the extended-open conformation, whereas 67% of the molecules in the presence of cilengitide and 60% of the molecules in the presence of MK-429 were in this conformation. None of the molecules in the presence of TDI-4161 were judged to have this conformation and only 5% of those in the presence of TDI-3761 had this conformation, again similar to the control sample. Together, these data provide graphic support for the hypothesis that TDI-4161 and TDI-3761 are pure αVβ3 antagonists. They also support the use of the AP5 screening assay to identify such compounds.

**Fig. 4.**
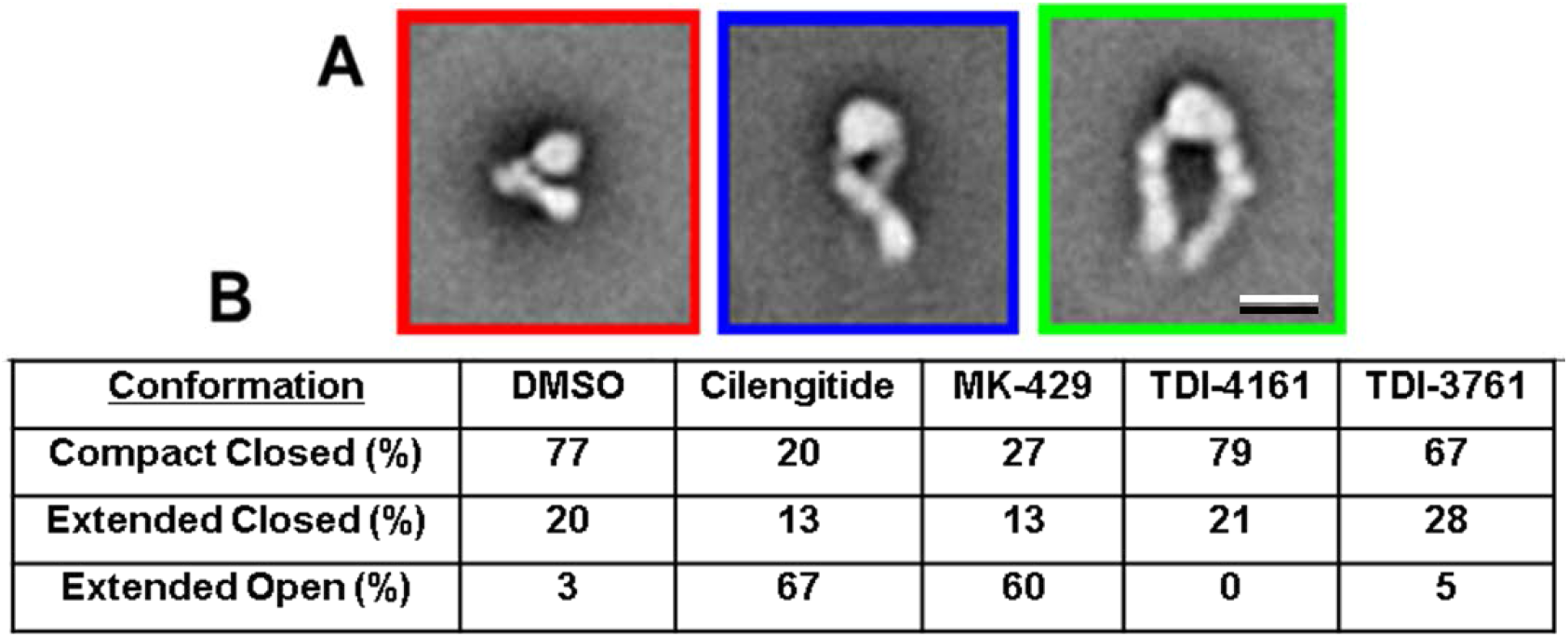
EM analysis of the effect of αVβ3 antagonists on αVβ3 conformation. (A) Typical averages of classes categorized as being in the compact-closed (red border), extended-closed (blue border), or extended-open (green border) conformation. Scale bar = 10 nm. (B) Percentage of molecules in each of the conformational states in the presence of DMSO or one of the αVβ3 antagonists (all at 10 μM).

### Specificity of TDI-4161 and TDI-3761

#### Selectivity for αVβ3 over αIIbβ3

We tested TDI-4161 and TDI-3761 for their ability to inhibit adhesion of HEK-293 cells expressing αIIbβ3 to fibrinogen (Fig. S8). Even at 10 μM, neither compound inhibited adhesion, which is similar to the control sample without any addition, or the vehicle control with 0.1% DMSO, whereas all of the positive control compounds [EDTA (10 mM), mAbs 7E3 and 10E5 (both at 20 μg/ml), and tirofiban (2 μM)] produced nearly complete inhibition. mAb LM609, which inhibits αVβ3 but does not react with αIIbβ3, also did not inhibit the adhesion of the cells expressing αIIbβ3 to fibrinogen.

#### Inhibition of ligand binding to purified αVβ3 and αVβ5

Cilengitide, MK-429 racemate, TDI-4161, and TDI-3761 all inhibited the binding of purified human αVβ3 to adenovirus 2 penton base with IC_50_s similar to those derived from studies of the adhesion of HEK-293 cells expressing human αVβ3 to fibrinogen (Table 1). With cilengitide and MK-429 racemate, the IC_50_s for the binding of purified αVβ5 to vitronectin were similar to those for inhibition of αVβ3-mediated cell binding to fibrinogen and purified αVβ3 binding to the penton base; in contrast, the IC_50_s for TDI-3761 (0.875 ± 0.348 μM) and TDI-4161 (0.689 ± 0.374 μM) were considerably higher than for αVβ3-mediated cell binding to fibrinogen or purified αVβ3 binding to penton base.

### Reactivity with mouse αVβ3

As a prelude to conducting studies in mice, we assessed the ability of the αVβ3 compounds to inhibit mouse αVβ3 receptors on endothelial cells. Table 1 shows the results in which the compounds were tested for their ability to inhibit the adhesion of mouse endothelial cells expressing αVβ3 receptors to immobilized fibrinogen. The MK-429 racemate had the lowest IC_50_, followed by cilengitide, TDI-3761, and TDI-4161.

### Priming studies

The results of the studies in which TDI-3761 and TDI-4161 were tested for their ability to prime the αVβ3 receptor to bind fibrinogen spontaneously are shown in Fig. 5. The RGDS peptide and cilengitide both significantly increased the binding of fibrinogen (p<0.001 and p<0.002, respectively, n = 10). In contrast, neither TDI-4161 (n = 10) nor TDI-3761 (n = 3) increased the binding of fibrinogen above the control level and each of their values was significantly lower than the value for cilengitide (p<0.002 and p<0.02, respectively).

**Fig. 5.**
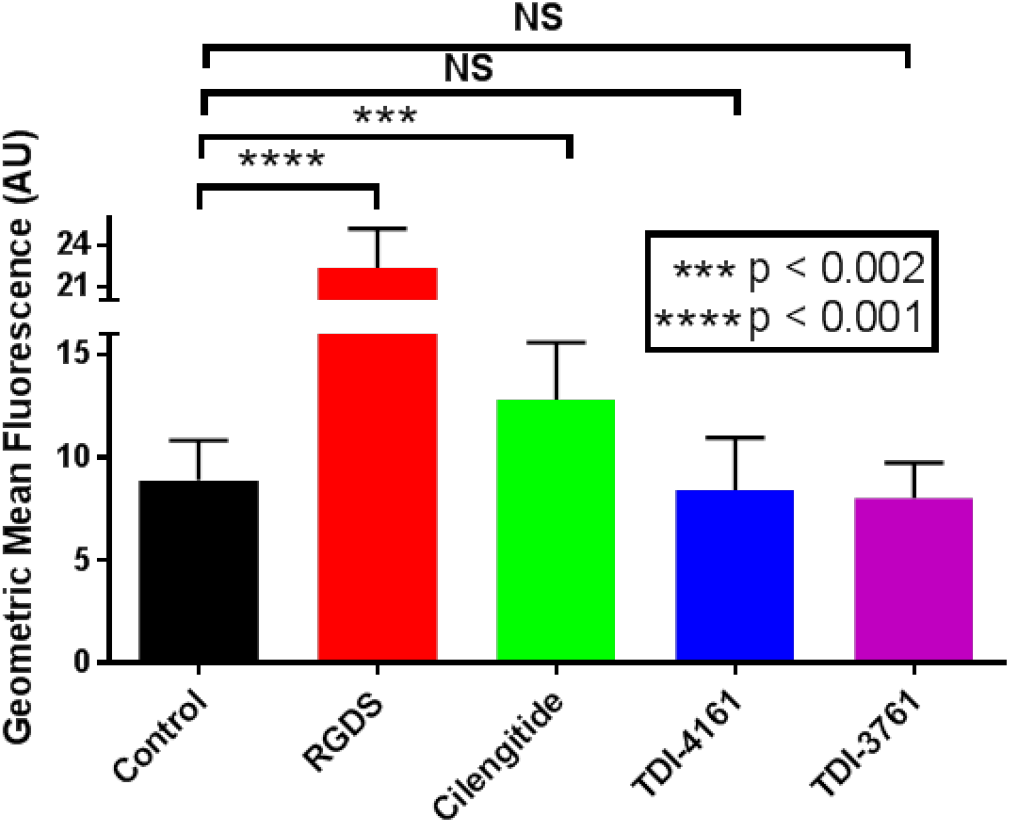
Priming of αVβ3. HEK-αVβ3 cells were either untreated (control) or incubated with 1 μM cilengitide, 100 μM RGDS, or 10 μM TDI-4161 or TDI-3761 for 20 minutes at room temperature, fixed with paraformaldehyde, washed, and incubated with fluorescent fibrinogen. After washing, cell-bound fluorescence was determined by flow cytometry. Compared to the control, both RGDS and cilengitide increased the amount of bound fibrinogen, whereas TDI-4161 and TDI-3761 did not. N = 7 for all values except TDI-3761, where n = 3.

### Osteoclast studies

#### Differentiation of murine bone marrow macrophages into osteoclast-like cells in plastic microtiter wells

When murine bone marrow macrophages in plastic microtiter plates were grown in the presence of RANK ligand, a source of M-CSF, and 0.1% DMSO for 3 days, they developed into osteoclast-like polykaryon cells as judged by their expression of osteoclast-related gene products NFATc1, cathepsin K, and integrin subunit β3. Murine macrophages grown for the same amount of time without RANK ligand, but with a source of M-CSF, did not express these gene products (Fig. 6A). In separate experiments, murine macrophages grown in the presence of RANK ligand and a source of M-CSF developed osteoclast-like morphology and stained positive for the osteoclast marker tartrate-resistant acid phosphatase (DMSO control, Fig. 6B,C). A control compound of similar structure to TDI-4161 and TDI-3761 that did not inhibit the adhesion of cells expressing αVβ3 to fibrinogen did not affect osteoclast-like gene expression or morphology when added to the cultures on day 0 (Fig. 6B) or day 4 (Fig. 6C) at 10 μM final concentration. MK-429 racemate and cilengitide (both at 10 μM) did not affect osteoclast-like cell differentiation as judged by gene expression (Fig. 6A), but they both resulted in a marked diminution in the development of typical osteoclast morphology when added on either day 0 or day 4 (Fig. 6B,C). Both TDI-4161 and TDI-3761 at 10 μM final concentrations also did not affect osteoclast-like cell differentiation as judged by gene expression (Fig. 6A). They did decrease the percentage of cells showing the typical osteoclast morphology (Fig. 6B,C), but not to the same extent as with either MK-429 racemate or cilengitide.

**Fig. 6.**
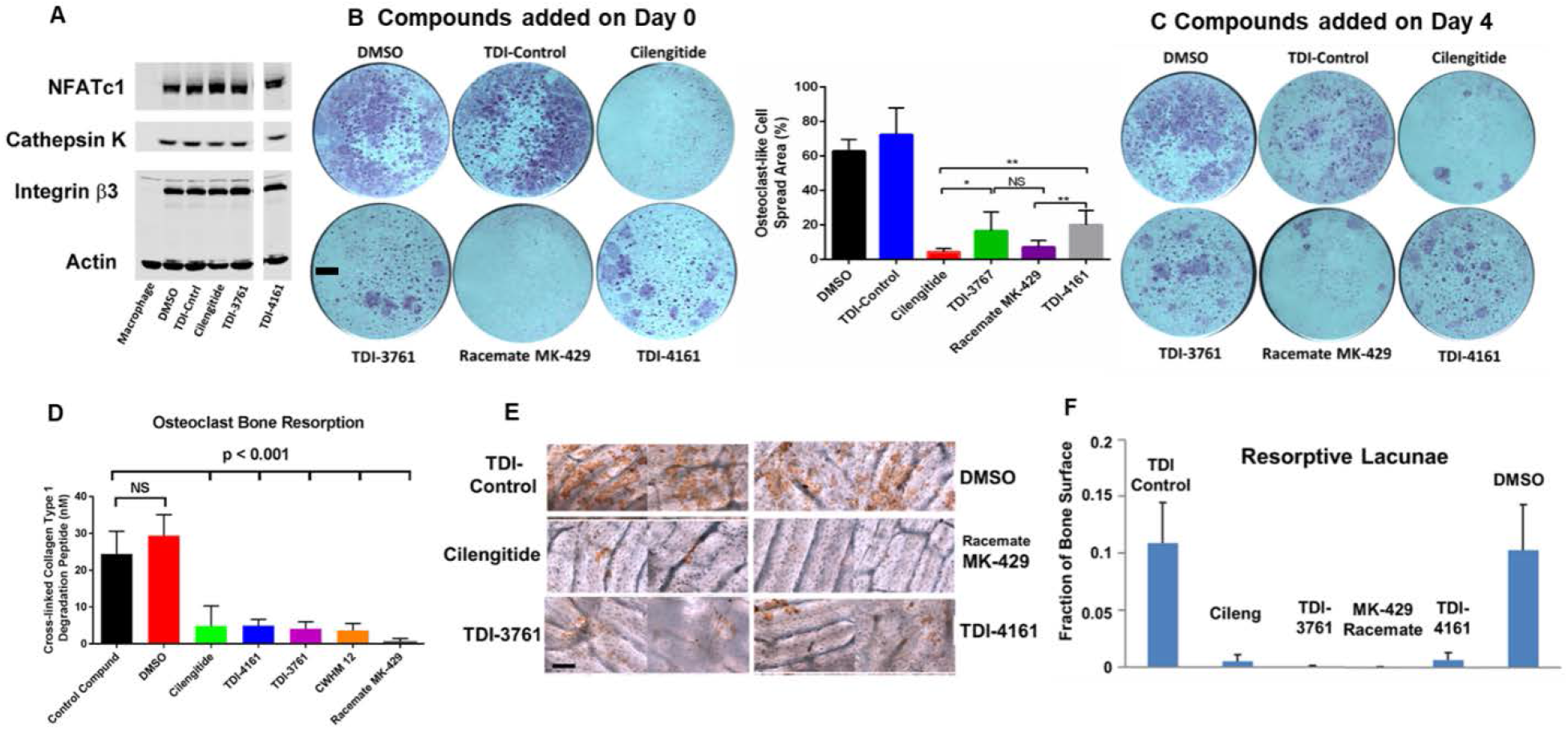
Effect of compounds on differentiation of murine bone marrow macrophages into osteoclast-like cells in the presence of RANK ligand and a source of M-CSF. (A) Immunoblot of expression of osteoclast marker proteins by murine macrophages after 3 days of culture in plastic wells in the presence of RANK ligand and a source of M-CSF. (B and C) Morphology of cells grown for 5 days on plastic and stained for the osteoclast marker tartrate resistant acid phosphatase. Compounds were added either on day 0 (B, top) or day 4 (C). Scale bar = 1 mm. Image analysis of the day 0 data is shown in the bottom panel of (B) (mean ± SD; Student’s *t*-test; n = 6 for each compound, except cilengitide and TDI-4161, where n = 5). (D) Resorption of bone by osteoclast-like cells as reflected in release of cross-linked collagen type 1 degradation peptide. Scale bar = 100 μm. (E) Direct staining of resorption lacunae (brown reaction product) produced on bone. (F) Quantification of fraction of the bone area with resorption lacunae [all compounds p<0.001 compared to DMSO Control except TDI Control (p=0.99) by ANOVA analysis with Dunnett post hoc test with adjustment for multiple testing].

#### Release of cross-linked collagen type 1 telopeptides by osteoclast-like cells on bovine bone

The concentrations of cross-linked collagen telopeptides, which are surrogate indicators of bone resorption,^69^ released into the medium collected on day 6 when osteoclast-like cells were grown on bovine bone are shown in Fig. 6D. The values for cells grown in the presence of DMSO or the control TDI compound, 29.4 ± 5.7 and 24.4 ± 6.2 nM, respectively (mean ± SD of quintuplicate replicates of a single experiment), were not significantly different (p = 0.22), whereas the values for all of the samples grown in the presence of the other compounds were significantly reduced relative to the control compound (p < 0.001 for all), with MK-429 racemate producing the greatest inhibition of release.

#### Lacunae formation on bovine bone slices

When grown from day 0 in the presence of DMSO or a control compound the osteoclasts produced bone resorption lacunae in the bovine bone that occupied 10.3 ± 4.0 and 10.9 ± 3.5% (mean ± SD of quintuplicate replicates of a single experiment) of the bone surface, respectively (Fig. 6E,F). In contrast, when the studies were performed in the presence of cilengitide or the racemate of MK-429, lacunae formation was reduced by more than 90% (to 0.53 ± 0.56 and 0.07 ± 0.03%, respectively; p < 0.001 for both *versus* DMSO or control compound). TDI-3761 and TDI-4161 also both inhibited osteoclast-mediated bone resorption lacunae formation, with values of 0.07 ± 0.06 and 0.68 ± 0.61% (p<0.001 for both versus DMSO or control compound).

### De-adhesion studies

The osteoclast-like cell studies indicated that MK-429 racemate, cilengitide, TDI-4161, and TDI-3761 had similar effects on cell morphology when they were added at the beginning of the culture or just on the last day before harvest, suggesting that they could not only prevent the cells from developing the typical morphology, but could actually reverse the morphology once it was established. To study this finding in more detail, we developed an assay in which αVβ3-expressing HEK-293 cells were allowed to adhere to fibrinogen for 30 minutes and were then treated with compounds and monitored over the next 30-60 minutes for de-adhesion as judged by the number of cells that remained adherent after additional washing. Adding MK-429 racemate, cilengitide, TDI-4161, or TDI-3761 resulted in profound inhibition of adhesion when added before adhesion, and equally profound de-adhesion after 30 minutes when added after the cells were adherent (Fig. S9A, Videos S1-S5; p<0.001 for all compounds compared to untreated cells, DMSO-treated cells, and cells treated with the TDI control compound).

We then assessed whether the AP5 epitope was exposed on the de-adherent cells by incubating AP5 with the cells that eluted from the fibrinogen-coated surface with washing (MK-429 racemate, TDI-4161, TCI-3761, and cilengitide) or cells that had to be removed from the fibrinogen-coated surface by trypsin treatment (control compound and DMSO) because they did not elute with washing. The AP5 epitope was exposed at between ~60% and ~50% of maximal on cells treated with MK-429 racemate and cilengitide in solution, respectively, and cells eluted from fibrinogen by these compounds showed only slightly lower levels of AP5 exposure (Fig. S9B; p<0.001 compared to control compound for all values). The cells treated with TDI-4161 or TDI-3761 in solution showed minimal AP5 exposure compared to control compound or the DMSO control, and similarly, the cells eluted from fibrinogen by these compounds demonstrated minimal AP5 binding. The cells treated with DMSO or the control compound in solution bound minimal amounts of AP5, as did the cells removed from the fibrinogen-coated surface by trypsin. We conclude that TDI-4161 and TDI-3761 can cause de-adhesion of αVβ3-expressing cells from fibrinogen without inducing the conformational change in β3 that exposes the AP5 epitope.

### Enhancement and inhibition of Vascular Endothelial Growth Factor (VEGF)-induced aortic ring endothelial cell sprouting

The outgrowth of microvascular sprouts from explanted mouse aortic rings can be used to test the efficacy of pro- and anti-angiogenic agents.^57,70^ The DMSO vehicle for the compounds did not modify the basal induction of sprouting (Fig. 7). VEGF is a potent pro-angiogenic factor in this assay, resulting in increased numbers of endothelial sprouts.^57,70^ Here we show that compared to the VEGF + DMSO vehicle control sample, cilengitide at 1 and 10 nM significantly enhanced VEGF-stimulated sprout formation by 43 and 24% (p=0.03 and p=0.04), respectively, while 100 nM and 10 μM inhibited sprout formation by 38 and 33% (p=0.004 and p=0.02, respectively).^57^ TDI-4161 at 1, 10, and 100 nM had no significant effect on VEGF-induced sprouting, while at 1 μM and 10 μM it enhanced sprouting by 45% and 50% (p=0.02 and p=0.03, respectively). TDI-3761 at 1, 10, and 100 nM had no significant effect on sprout formation; at 1 μM it inhibited sprouting by 26% (p=0.03), and at 10 μM it enhanced sprout formation by 36%, but the result was not significant (p=0.14). We conclude that: (1) while cilengitide enhances VEGF-induced angiogenesis at 1 and 10 nM, concentrations below its IC_50_ for inhibiting αVβ3-mediated murine cell adhesion (26 nM), TDI-4161 demonstrates enhancement of angiogenesis only at 1 and 10 μM, concentrations much above its IC_50_ (67 nM), and TDI-3761 shows only a trend toward enhancing angiogenesis at 10 μM, which is also much above its IC_50_ (41 nM), and (2) cilengitide significantly inhibits angiogenesis at concentrations above its IC_50_ (>100 nM), whereas only TDI-3761 at 1 μM significantly inhibits angiogenesis.

**Fig. 7.**
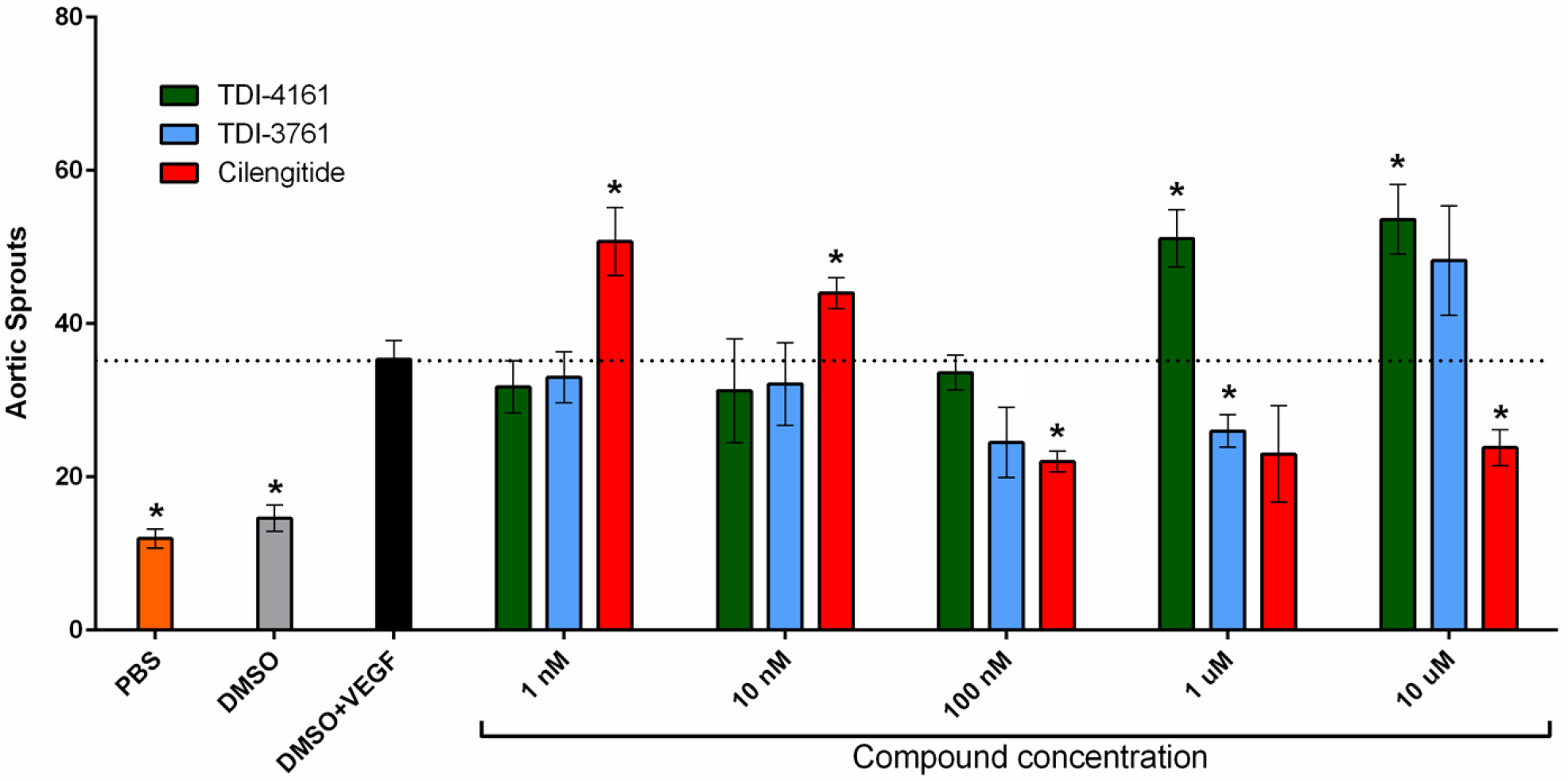
Effect of αVβ3 targeting compounds on VEGF-induced angiogenesis *ex vivo*. (A) Mouse aortic rings were stimulated with OptimMEM supplemented with 2.5% FCS and 30 ng/ml VEGF, PBS, or vehicle alone (DMSO). Cilengitide, TDI-4161, or TDI-3761 were added to OptimMEM supplemented with 2.5% FCS and 30 ng/ml VEGF at 1 nM, 10 nM, 100 nM, 1 μM and 10 μM. The number of sprouts per ring was counted in a blinded fashion using a phase contrast microscope at day 8 post embedding. Data are presented as mean ± SEM. To minimize the potential impact of inter-animal variations in angiogenesis, rings from the aortas of four animals were included in each experimental condition. The number of rings included in each experimental condition varied, however, from one to four. To prevent overweighting the impact of any aorta, the number of sprouts from the rings from the same aorta in each experimental condition were averaged, yielding four values for each condition, one for each aorta. The data presented are the mean ± SEMs of these four values. Asterisks above the data bars indicate that the results differ significantly (p<0.05) from that of the DMSO + VEGF sample using two-tailed Student’s *t*- test without correction for multiple comparisons.

## Discussion

There are many potential therapeutic applications of αVβ3 antagonists, but currently none is approved for human use. The failure of cilengitide to demonstrate efficacy in treating glioblastoma^34^ may be due to differences in the pathophysiology of the human disease and the animal models in which it showed efficacy. An alternative possibility is that cilengitide’s antitumor effects are limited by its paradoxical effects in enhancing tumor growth and tumor angiogenesis in animals at low concentrations, which we previously demonstrated correlates with increased endothelial cell cycling of VEGFR2.^57^ Since these effects did not occur in mice lacking β3,^57,71^ it is likely that they are due to cilengitide engaging endothelial cell αVβ3 and initiating signaling through the receptor. Our current data showing that cilengitide has a nanomolar EC_50_ for AP5 exposure are consistent with the hypothesis that cilengitide’s ability to initiate the conformational change in αVβ3 underlies its ability to enhance VEGFR2 cycling. One possible reason that the effect is only observed at sub-saturating concentrations of cilengitide is that after binding to αVβ3 and inducing signaling and adoption of the high-affinity ligand-binding state, it releases from the receptor and leaves it unoccupied and in the high-affinity ligand-binding state.^72^

A similar mechanism may also limit the ability of partial agonists to treat renal podocyte disorders^18^ and supravalvular aortic stenosis in William’s syndrome^29^ since signaling through αVβ3 may contribute to both of these disorders. Thus, it would be valuable to have high potency pure αVβ3 antagonists to assess whether they offer insights into the pathophysiology of the disorders and/or therapeutic advantages.

Our studies demonstrate that it is possible to exploit the structural information obtained by Arnaout’s group on the interaction of the pure peptide antagonist hFN10 that interacts with the RGD pocket in αVβ3 but does not induce the conformational change in the β3 subunit associated with the receptor adopting a high-affinity ligand-binding state^62^ to design small molecules with similar properties. Guided by the predicted binding mode of the high-affinity αVβ3 antagonist MK-429, we simplified the molecule and then extended its length and substituted the α carbon to position a bulky aromatic group adjacent to β3 Tyr122 (the same position occupied by Trp1496 of hFN10) in an attempt to prevent the latter’s movement toward the MIDAS when the carboxyl group engages the MIDAS and interacts with the backbone nitrogens in the β3 β1-α1 loop. Among the compounds that we synthesized, TDI-3761 and TDI-4161 have nanomolar IC_50_s, but unlike MK-429 and cilengitide, αVβ3 antagonists patterned on the RGD sequence, these compounds do not expose the epitope for AP5 or prime the receptor to bind ligand even at 10 μM.

We previously demonstrated using negative-stain EM that cilengitide could induce a dramatic change in the conformation of αVβ3, converting it from a compact-closed conformation to an extended-open conformation that could also be induced by the integrin-activating agent Mn^2+^^58^. Thus, cilengitide induced the receptor to adopt the high-affinity ligand-binding conformation, demonstrating its activity as a partial agonist. Our current EM studies extend these observations to demonstrate that MK-429 has the same partial agonist activity, whereas TDI-4161 and TDI-3761 have little or no ability to induce the conformational change. This result provides strong support for the hypothesis that TDI-4161 and TDI-3761 are pure antagonists and thus supports our choice of using AP5 epitope exposure as a screening strategy for classifying antagonists.

The crystal structure of the human αVβ3 ectodomain in complex with TDI-4161 showed that the βA domain assumed the inactive conformation despite the carboxyl oxygen coordinating the MIDAS Mn^2+^, and we conclude that this is due to *π*-*π* stacking of the benzothiazole group of TDI-4161 against β3-Tyr122, in accord with the results from docking and MD studies. This interaction cannot be formed by the *R*-enantiomer of TDI-4161, offering an explanation for the difference in potency of the enantiomers. The specificity of TDI-4161 for αVβ3 but not αIIbβ3 is explained by the inability of the Arg-mimetic naphthyridine group to make a salt bridge with αIIb Asp224, which is required for binding to αIIb. The direct involvement of β3-Arg214 (present only in β3 and β5) and Met180 (unique to β3) in binding TDI-4161 is consistent with its higher affinity for αVβ3 versus αVβ5. Human β3-Tyr122 is replaced with Phe122 in mouse β3, and the stabilizing salt bridge β3-Arg214 makes with β3-Asp179 is replaced with an H-bond with Asn179 in mouse β3, both substitutions likely contributing to the observed lower affinity of TDI-4161 for mouse αVβ3.

The Gln319-Ser674 contact between the βA and βTD domains is absent in structures of unliganded αVβ3^73^ or αVβ3 in complex with natural ligands^62^ or partial agonists like cilengitide,^54^ suggesting that separation of these two domains is an important component of the conformational pathway leading to αVβ3 activation.

TDI-3761 and TDI-4161 both inhibited murine αVβ3-mediated osteoclast-like cell spreading and bone resorption *in vitro*, demonstrating αVβ3 target engagement and integrin antagonism. Of note, both compounds, along with the racemate of MK-429 and cilengitide, reversed the spread morphology when added after the murine bone marrow macrophages differentiated into osteoclast-like cells, suggesting that the compounds could cause the αVβ3 receptors to disengage from ligand. To further assess this possibility, we developed a de-adhesion assay and found that the compounds could cause cells adherent to fibrinogen via αVβ3 to de-adhere when exposed to the compounds. Even after de-adhesion, the αVβ3 receptors on the cells treated with TDI-3761 and TDI-4161 did not expose the AP5 epitope, whereas those treated with MK-429 racemate and cilengitide did expose the AP5 epitope.

The demonstration of a compound’s ability to disengage αVβ3 from ligand is particularly important because, unlike the unoccupied αIIbβ3 receptors on resting platelets, αVβ3 receptors implicated in the pathogenesis of disease are presumed to be engaged with extracellular matrix proteins under basal conditions, and in some disorders, the activation of the receptor has been documented by the ability of AP5 or the ligand-mimetic mAb WOW-1^74^ to bind to the receptor in the affected tissue.^29,75,76^ In fact, αVβ3 receptors on different cell types appear to have different levels of both basal activation and responsiveness to activation by different agents.^74^

As indicated above, high nanomolar concentrations of cilengitide may inhibit tumor growth, whereas low nanomolar concentrations may have an opposing effect, providing one potential explanation for the lack of efficacy of cilengitide in the treatment of glioblastoma.^57^ It is of particular interest, therefore, that TDI-4161only demonstrated enhancing effects on aortic sprout formation at micromolar concentrations and TDI-3761 showed only a trend toward enhanced sprout formation at 10 μM. Since these compounds inhibit αVβ3-mediated cell adhesion at nanomolar concentrations, it may be possible to develop a dosing strategy that will maintain effective αVβ3 receptor inhibition without enhancement of angiogenesis and related phenomena. Whether the enhanced angiogenesis at high concentrations of TDI-4161 and the similar trend with TDI-3761 represent a small residual ability to initiate a conformational change at very high doses or some other effect remains to be studied. Similarly, whether these effects can be eliminated by further medicinal chemistry structural refinements also remains to be studied.

On the other hand, TDI-4161 and TDI-3761 were less effective at inhibiting angiogenesis than cilengitide, with only TDI-3761 demonstrating significant inhibition at 1 μM. This may be due to cilengitide’s inhibition of αVβ5 in addition to αVβ3 since αVβ5 has also been implicated in contributing to angiogenesis;^77^ TDI-4161 and TDI-3761 are more specific for αVβ3. Thus, whether TDI-4161 and TDI-3761’s differences in inducing and inhibiting angiogenesis from cilengitide will translate into improved therapy of malignancy remains to be established in additional animal models and human studies.

One way to assess the likely potential toxicities of long-term inhibition of αVβ3 is to analyze the phenotypes of patients who lack αVβ3 on a genetic basis. Patients with defects in the integrin β3 subunit lack both αVβ3 and the integrin receptor αIIbβ3, which is specific for platelets and megakaryocytes and plays a vital role in platelet aggregation. Thus, these patients have a life-long bleeding disorder (Glanzmann thrombasthenia).^78^ There have been no specific medically significant defects identified in these patients beyond their bleeding diathesis, although relatively few patients have been studied in detail.^79^ This finding is consistent with the generally favorable safety profile for the pan-αV receptor antagonist MK-429^80^ when administered to 116 postmenopausal women with osteoporosis for 12 months at varying doses.^36^

Based on the evidence that αVβ3 plays an important role in bone resorption, it is possible that chronic therapy in non-osteoporotic patients may result in abnormally high bone mineral density. To assess this possibility we previously studied bone mineral density in a group of five female Glanzmann thrombasthenia patients age 39-57 with defects in β3^81^ and did not identify a consistent increase in bone density.^82^ This is paradoxical in view of: (1) The demonstrated age-dependent increase in bone density in mice lacking αVβ3 and with conditional targeting of myeloid cell β3,^83,84^ although the increase in bone density in these animals is modest,^83–85^ and (2) evidence that mice lacking αVβ3 are protected from developing loss of bone mineral density after ovariectomy.^86^ One possible explanation for the difference in the human and mouse phenotypes comes from the study by Horton et al. of osteoclast-like cells derived from the peripheral blood mononuclear cells of patients with a defect in β3.^87^ Unlike the severe abnormality in spreading and actin ring formation we and others have observed in cultured bone marrow macrophages from mice lacking β3,^83–85^ which correlate with their defect in bone resorption, they found that the osteoclast-like cells from the patients had relatively normal spreading and actin ring formation.^87^ Patient cells were less effective in resorbing bone, however, with a 44% decrease in the number of lacunae and a 59% decrease in the depth of the lacunae; these abnormalities are not as profound as we observed with our αVβ3 antagonists, but are similar to those reported in β3-null mice.^83^ They ascribed the relatively preserved patient osteoclast-like cell phenotype to a 2-4-fold increase in α2β1 expression facilitating interaction with collagen. Thus, it is possible that there is functional integrin or non-integrin receptor compensation in humans with hereditary loss of β3, leading to a milder phenotype. Alternatively, αVβ3 may play an enhanced role in osteoclast bone resorption in the post-menopausal state relative to its role under basal conditions.

In summary, our data demonstrate the ability to develop small-molecule pure antagonists of αVβ3, providing vital tool compounds for dissecting the effect of inducing the activating conformational change in the receptor in the many pathological processes in which αVβ3 has been implicated. If pure antagonists demonstrate benefits over partial agonists in model systems, they may be appropriate to consider for human therapy.

## Materials and Methods

**See Supplementary text for details on Materials and Methods described below.**

### HEK-293 cells expressing αVβ3 (HEK-αVβ3)

HEK-293 cells were transfected with the cDNA for αV using the pEF1/V5-His A vector and the cDNA for β3 using the pcDNA3.1 vector. Cells expressing αVβ3 were identified and stable cell lines were established by repetitive sorting. HEK-αVβ3 cells for assays were counted and adjusted to values appropriate for each assay.

### αVβ3-mediated cell adhesion to fibrinogen assay

Polystyrene 96-well microtiter plates (Costar, 3590) were precoated with 3.5 μg/ml of purified fibrinogen and incubated with HEPES-Buffered Modified Tyrode’s solution [HBMT; 0.128 M NaCl, 10 mM HEPES, 12 mM NaHCO_3_, 0.4 mM NaH_2_P0_4_, pH 7.4, 2.7 mM KCl, 0.35% bovine serum albumin (Fisher), 0.1% glucose] for 1 hour at room temperature or overnight at 4°C. Wells were washed with HBMT containing 1 mM Mg^2+^ and 2 mM Ca^2+^ and then 50 μl of HEK-αVβ3 cells that were pretreated with the compound to be tested for 20 minutes at room temperature were added to each well at a concentration of 3,000 cells/μl. After ~30 minutes the wells were washed three times with HBMT containing Ca^2+^ and Mg^2+^ and then the adherent cells were lysed and the acid phosphatase activity that was released was measured. In each assay, 10 mM EDTA was used as a positive control and untreated cells were used as a negative control. The IC_50_ was defined as the concentration of the test compound that reduced the adhesion of the HEK-αVβ3 cells by 50%, taking the results with untreated cells as 100% and the results in the presence of EDTA as 0%.

### AP5 binding assay

HEK-αVβ3 cells were harvested, washed with HBMT once, and resuspended in HBMT containing 1 mM Mg^2+^ and 2 mM Ca^2+^. 5 × 10^5^ cells were incubated with fluorescently labeled mAb AP5 for 30 minutes at 37°C. The cells were then washed and analyzed by flow cytometry. In each assay, cilengitide (1 μM) and 10 mM EDTA were included as positive controls and untreated cells were used as the negative control. The concentration of the test compound required to induce half-maximal exposure of the AP5 epitope as judged by exposure produced by 1 μM cilengitide was calculated and defined as the EC_50_. The EC_50_ for cilengitide was determined based on the exposure induced by EDTA. The AP5 exposure induced by 1 μM cilengitide was approximately twice the value with 10 mM EDTA [average ± SD of 17 experiments; control 7.6 ± 2.2, EDTA 21.8 ± 5.9, cilengitide 42.5 ± 8.0 arbitrary fluorescence units (AFU)]. In cases in which even the highest concentration of test compound (10 μM) did not induce 50% exposure of the AP5 epitope, the results are reported as >10 μM.

### Priming assay

HEK-αVβ3 cells were washed, resuspended in HBMT containing 1 mM Mg^2+^ and 2 mM Ca^2+^ at 2 × 10^6^/ml, and either left untreated (control) or incubated with 1 μM cilengitide, 100 μM RGDS, or 10 μM TDI-4161 or TDI-3761 for 20 minutes at room temperature. Samples were then fixed with 4% paraformaldehyde in phosphate-buffered saline (PBS) for 40 minutes at room temperature, followed by quenching of the reaction with 5 mM glycine for 5 minutes at room temperature. After washing with HBMT, the cells were resuspended in HBMT containing 1 mM Mg^2+^ and 2 mM Ca^2+^. Alexa488-conjugated fibrinogen was then added and incubated for 30 minutes at 37°C. The cells were then washed and analyzed by flow cytometry.

### De-adhesion assays

We modified the assay of Charo et al. for human endothelial cells.^88^ Adhesion of HEK-293-αVβ3 cells to fibrinogen was carried out as above for 30 minutes in the absence of compounds and unattached cells were removed by washing. Compounds were then added and after an additional 30-60 minutes, the wells were washed again and the number of remaining cells was analyzed and compared to the number of cells that adhered during the first 30 minutes. In some experiments, AP5 binding was performed on cells that de-adhered during the experiment in the presence of compound. Since cells in the control sample did not de-adhere during the additional 30-60 minutes, they were lifted from the plate by treatment with trypsin, but without EDTA. For time-lapse studies of the de-adhesion process, HEK-αVβ3 cells (2 × 10^6^/ml) were plated on IbiTreat μ-Slide 8 wells (ibidi, Martinsried, GmbH) pre-coated with fibrinogen. Differential Interference Contrast (DIC) images were acquired using a water immersion objective. Final movies were processed and assembled using FIJI/Image J. Channels were gamma-adjusted to enhance visualization.

### Mouse endothelial cell adhesion assay

Mouse primary aortic endothelial cells were grown in endothelial cell medium (M1168), harvested with trypsin-EDTA, washed once with HBMT, and resuspended in HBMT containing 1 mM Mg^2+^ and 2 mM Ca^2+^. The cells were then incubated with the test compounds for 20 minutes at 22°C and added to microtiter wells that had been pre-coated by adding 50 μl of human fibrinogen (5 μg/ml) in Tris-saline buffer, pH 7.4 at 4°C overnight and then washing and blocking with HBMT containing 0.35% albumin. The cells were allowed to adhere for 30 minutes at 37°C, after which the wells were washed with HBMT containing Mg^2+^ and Ca^2+^ and the number of remaining adherent cells was assessed by lysing the cells in Triton X-100 and determining the acid phosphatase activity.

### Inhibition of ligand binding of purified αVβ3 and αVβ5

Purified receptors were obtained from R&D Systems and tested by modifications of the assays described by Henderson et al.^20^. The ligands employed were adenovirus 2 penton base^89^ (kindly supplied by Dr. Glen Nemerow of Scripps Research Institute) for αVβ3 and vitronectin for αVβ5. Microtiter wells were coated overnight at 4°C, and then washed with TTBS buffer (137 mM NaCl, 25 mM Tris/HCl, pH 7.4, 2.7 mM KCl, 0.1% Tween-20) and blocked with 1% bovine serum albumin (BSA) for 60 minutes at 22°C. The purified receptor in TTBS + 0.1% BSA was incubated with the compound to be tested for 20 minutes at 22°C and then the mixture was added to the well. After 120 minutes at 37°C, the wells were washed with TTBS and the bound receptor measured by adding a biotin-labeled mAb to αV and detecting the antibody by using horse radish peroxidase-labeled streptavidin. After subtracting the value obtained in the presence of EDTA (30 mM) from each result, the percentage inhibition of binding and the IC_50_ for each compound was determined as above.

### Inhibition of αIIbβ3-mediated cell adhesion to immobilized fibrinogen

The ability of the compounds to inhibit the interaction between αIIbβ3 and fibrinogen was measured by assessing their ability to inhibit the binding of HEK-293 cells expressing αIIbβ3 to immobilized fibrinogen as reported previously.^47^

### Organic synthesis of select compounds

The details of the synthesis of each of the compounds is provided in Supporting Information.

### Osteoclast culture for morphology, bone lacunae formation, and cross-linked collagen degradation peptides

All studies were performed on coded samples with the experimenter not knowing the identity of the individual compounds. As previously described,^85^ bone marrow was harvested from the femur and tibia of 8-10 week old male C57BL mice and cultured in α modification of Minimal Essential Medium (α-MEM) supplemented with 10% fetal bovine serum and 10% conditioned medium from the CMG14-12 cell line^90^ as a source of M-CSF for 4 days. The resulting macrophages were collected with trypsin-EDTA and cultured in plastic tissue culture plates with or without bovine bone slices (1.2 × 10^4^ cells/well/0.5 ml medium) with RANK ligand (100 ng/ml), M-CSF (2% CMG-conditioned medium), test compounds, or DMSO (1:1000 final dilution). In some experiments, compounds were added on day 4 instead of on day 0.

On day 5, cells grown in the plastic plate with bone were fixed with 4% paraformaldehyde for 10 minutes at room temperature and stained for tartrate-resistance acid phosphatase. The surface spread area percentage was quantified using Fiji software.^91^ The sum of all of the surface areas per well was calculated and converted into percentage using the area of an empty well as 100%.

On day 6, the medium of cells grown on bone was collected for assay of cross-linked degradation products of collagen type 1 telopeptides. The cells grown on bone for osteoclast resorption lacunae analysis were removed from the bone slices on day 6 and the bone slices were incubated with peroxidase-conjugated wheat germ agglutinin and then stained with 3,3’- diamaninobenzidine. Five defined 10x fields were photographed and the percentage of the image containing lacunae was determined by image analysis. Each compound was studied on 5 separate bone slices.

In other experiments, murine bone marrow macrophages prepared as above were cultured with RANK ligand and M-CSF in the presence of DMSO or test compounds for 3 days and then cells were lysed and tested for osteoclast differentiation markers (integrin subunit β3, NFATc1, and cathepsin K) by immunoblotting. Bone marrow macrophages cultured in M-CSF alone served as a negative control.

### Mouse aortic ring vascular sprout assay

These studies were conducted in accord with the policies of the Ethics Committee of Queen Mary University of London. 8 C57Bl6 mice were euthanized at 8-10 weeks of age and their thoracic aortae dissected. As previously described,^70^ aortae were cut into 0.5-mm rings, starved overnight in OptiMEM medium with penicillin-streptomycin, and then embedded in rat tail collagen type 1. Aortic rings were then stimulated with 150 μl of OptimMEM supplemented with 2.5% FCS and 30 ng/ml VEGF, PBS, or vehicle (DMSO) alone. Cilengitide, TDI-4161, or TDI-3761 was added to OptimMEM supplemented with 2.5% FCS and 30 ng/ml VEGF at 1 nM, 10 nM, 100 nM, 1 μM and 10 μM. The number of sprouts per ring was counted using a phase contrast microscope at day 8. After that, rings were fixed, permeabilized, blocked using 2% BSA, and stained with TRITC-conjugated lectin from *Bandeiraea simplicifolia*. Images were captured using a LSM710 confocal microscope.

A total of 8 aortas were used to prepare the rings for the study, and each experimental condition included rings derived from the aortas from four animals. To minimize the potential impact of inter-animal variations in angiogenesis, the number of sprouts observed in each ring derived from a single aorta under each experimental condition were averaged. The primary hypothesis was that compounds TDI-4161 and TDI-3761 would not enhance VEGF-induced sprout formation at sub-IC_50_ concentrations, whereas based on our previous results, cilengitide would enhance sprout formation at sub-IC_50_ concentrations and inhibit sprout formation at higher concentrations. As a result, we compared the results of the DMSO + VEGF sample to those of TDI-4161 and TDI-3761 using a two-sided t test without correction for multiple comparisons.

### Molecular docking

After removal of the high-affinity recombinant fibronectin hFN10 domain, the crystal structure of αVβ3 corresponding to PDB code 4MMZ was used to dock either the potent αVβ3 inhibitor MK-429, the lead compound TDI-4161, or the *S*-enantiomer of the racemate TDI-3761, with the Glide v6.6 docking algorithm included in the Schrödinger Suite 2015-1. The receptor was prepared using Maestro v10.1 while the ligands were prepared with LigPrep v3.3. The nitrogens on the naphthyridine moiety were protonated to mimic the bidentate interaction formed by Arg1493 of the fibronectin RGD motif with the αV residue Asp218 as seen in the 4MMZ crystal structure, whereas a charged carboxyl terminal mimicked the interaction between the Asp1495 sidechain of the fibronectin RGD motif and the metal ion in the MIDAS as also seen in the 4MMZ crystal structure. A grid box with outer and inner dimensions of 29 Å × 31 Å × 29 Å and 10 Å × 12 Å × 10 Å, respectively, was centered at the equivalent position of the fibrinogen R^1493^GDW^1496^ sequence in the 4MMZ crystal structure, and used for an initial Glide Single Precision (SP) docking followed by a Glide Extra Precision (XP) refinement.

### Molecular dynamics simulations

The αVβ3 receptor complexes with the top-scoring docking poses of MK-429 or TDI-4161 were subjected to standard MD simulations in an explicit solvent environment using the GROMACS simulation package and the CHARMM General Force Field (CGenFF) parameters for the ligands. To contain computation time, the simulated receptor system was limited to the head domain (αV residues 1-437 and β3 residues 109-352), which is most relevant to ligand binding. The ligand-receptor complex was solvated in a dodecahedron water box with a 7 Å minimum distance between the solute and the box edges, and counter ions were added to neutralize the system. The TIP3P water model and CHARMM36 force field were applied. Since parameters for the Mn^2+^ ions in the structure are not available in CHARMM36, Mg^2+^ ions were simulated instead. The final ligand‒receptor complex system contained the solute, ~24,000 water molecules, 7 Mg^2+^ ions, and 11 Na^+^ ions, totaling ~84,000 atoms.

Simulations were carried out with GROMACS v5.1.2. Prior to MD equilibration and production runs, energy minimizations were carried out using the steepest descent algorithm for 2000 steps. System equilibration consisted of 10 ps in the NVT ensemble with all solute heavy atoms constrained, and was followed by relaxations of 5 ns in the NPT ensemble with decreasing positional restraints, first on all solute heavy atoms and then on the protein Cα atoms and ligand polar atoms only. All restraints were removed prior to the production run, and the atomic velocities were randomized according to the Maxwell distribution at 300 K. Three independent production runs of at least 20 ns were performed in the NPT ensemble at 300 K and 1 bar using a V-rescale thermostat, Parrinello-Rahman pressure coupling, and a time step of 2 fs. All bonds were restrained using the LINCS algorithm, and a 12-Å cutoff was used for short-range non-bonded interactions. The root-mean-square deviations (RMSDs) of the ligand heavy atoms during the three independent MD runs of MK-429 or TDI-4161 bound to the protein are shown in Figs. S1 and S2, respectively. The RMSD was calculated after fitting protein heavy atoms onto the starting (docked) structure. Simulation trajectories were also examined for ligand-receptor interactions seen in the docking poses, which were plotted as center of mass (COM) distances in Figs. S3 and S4.

### Integrin expression, purification, and crystallography

Human αVβ3 ectodomain was expressed in insect cells, purified and crystallized at 4°C by the hanging drop method as previously described ^62^. TDI-4161 was soaked into the preformed αVβ3 crystals at 1 mM in 10% DMSO (v/v) in the crystallization well solution containing 1 mM Mn^2+^ for 3 days. Crystals were harvested in 12% PEG 4000 (polyethylene glycol, molecular weight 4000), 0.8 M sodium chloride, 0.1 M sodium acetate (pH 4.5), 1 mM Mn^2+^; cryoprotected with additional glycerol in 2% increments up to a 24% final concentration; and then flash-frozen in liquid nitrogen. Diffraction data were collected at ID-19 of the Advanced Photon Source (APS), indexed, integrated and scaled by HKL2000,^92^ and solved by molecular replacement using 3IJE as the search model in PHASER.^93^ The structure was refined with Phenix,^94^ using translation-liberation-screw, automatic optimization of X-ray and stereochemistry, and Ramachandran restriction in the final cycle. Data collection and refinement statistics are shown in Table S1. The coordinates and structure factors of the αVβ3-TDI-4161complex have been deposited in the Protein Data Bank under accession code 6MK0.

### EM sample preparation, imaging, and image processing

Recombinant αVβ3 ectodomain was produced and purified as previously reported,^95^ except that stably expressing HEK-293S GnT1- cells^96^ were used instead of CHO-lec cells. A 5-μL aliquot of integrin solution at 0.007 mg/ml was applied to a glow-discharged thin carbon film that was evaporated onto a plastic-coated copper grid. After 15 s, the grid was blotted, washed twice with deionized water and stained with 0.07% (w/v) uranyl formate as described.^97^ Grids were imaged with a Philips CM10 electron microscope operated at an acceleration voltage of 100 kV using a defocus of about –1.5 μm and a calibrated magnification of 41,513x, yielding a pixel size of 2.65 Å at the specimen level. For each of the 5 samples, 40 images were collected with an AMT 3K × 5K ActiveVu CCD camera. Gautomach (https://www.mrc-lmb.cam.ac.uk/kzhang/Gautomatch) was used to automatically and reference-free pick ~8,000 particles from 20 images of the cilengitide sample, and the particles were subjected to 2D classification in Relion.^98^ Three of the resulting class averages representing different αVβ3 conformations were then used to pick all the images of all 5 samples with Gautomach (the number of particles for each sample are listed in Fig. S7). The particles were extracted into 144×144-pixel images, centered and normalized in EMAN2.^98^ The particles were classified into 100 groups using *K*-means classification procedures implemented in SPIDER.^99^ Class averages with clear structural features were manually assigned to represent αVβ3 in a compact-closed, extended-closed, or extended-open conformation; the remaining averages were not assigned. The criteria for differentiating the extended-closed from the extended-open conformation included whether the legs are crossed and whether the β3 hybrid domain appears to face more toward the headpiece rather than out from the headpiece. The percentages provided in Fig. 4 were calculated as the fraction of integrins in a particular conformation with respect to all the assigned particles.

## Acknowledgments

We thank Dr. Peter Meinke for valuable advice and Suzanne Rivera for outstanding administrative support. This work was supported, in part, by grant HL19278 (B.S.C., M.F., Y.Z., and T.W.) from the Heart, Lung, and Blood Institute of the National Institute of Health; grants DK088327 (M.A.A.) and DK101628 (J.V.A.) from the National Institute of Diabetes and Digestive and Kidney Diseases of the National Institute of Health; UL1 TR001866 from the National Center for Advancing Translational Sciences of the National Institute of Health; the Tri-Institutional Therapeutic Discovery Institute (B.S.C. and M.F.); the Robertson Discovery Fund (B.S.C.); funds from Stony Brook University; grants 85400-STL from Shriners Hospitals for Children, AR046523 and AR057235 from the National Institute of Arthritis and Musculoskeletal and Skin Diseases, and DK111389 from the National Institute of Diabetes and Digestive and Kidney Diseases (S.L.T.); and funds from Worldwide Cancer Research (16-0390) (J.M-F.) and Cancer Research UK (C8218/A21453) (K.H-D). Computations were run on resources available through the Scientific Computing Facility at the Icahn School of Medicine at Mount Sinai and the Extreme Science and Engineering Discovery Environment under MCB080077 (M.F.), which is supported by National Science Foundation grant number ACI-1548562.

## Author contributions

J.L., L.B. and D.N. designed and conducted the IC_50_, EC_50_, and de-adhesion analyses. Y.F., R.H., Y.T., R.O., T.Y., T.N., T.I., K.A., C.L., M.D., and M.F. designed and performed the medicinal chemistry syntheses and characterized the resulting compounds. Y.S., Yu. Z., and M.F. designed, conducted, and interpreted the computational studies. J.M-F. and K.H-D. designed, conducted, and interpreted the aortic ring sprouting assays. W.Z. and S.T. designed, conducted, and analyzed the osteoclast-like cell studies. J.V.A. and M.A.A. expressed and purified recombinant human αVβ3 and determined and analyzed the crystal structure of the αVβ3-TDI-4161 complex. J.T. provided purified αVβ3 ectodomain for the EM studies. Yi. Z. and T.W. performed and interpreted the EM studies. R.V. performed the statistical analysis of the aortic sprout angiogenesis assay, B.C. conceived and oversaw the project, including recruiting collaborators, analyzing data, and taking primary responsibility for writing the manuscript.

