## Supplementary Materials for "Novel Pure αVβ3 Integrin Antagonists That Do Not Induce Receptor Extension, Prime the Receptor, or Enhance Angiogenesis at Low Concentrations"

**This PDF file includes:**

**Supplementary text**
- Materials and Methods
- Synthesis of individual compounds

**Supplementary figures and table**
- Figs. S1 to S9 and Table S1

**References for SI reference citations.**

**Supplementary movies**
- Movie S1-S5 - **Videos of de-adhesion studies**. HEK-αVβ3 cells (2 x $10^6$ cells/ml) were plated on an IbiTreat µ-Slide 8 well pre-coated with fibrinogen for 30 minutes. Differential interference contrast (DIC) images were acquired on a Leica SP8 inverted confocal microscope using a water immersion objective. For pre-treatment, DIC images were taken every 7 s for a 5-minute period. After this time, the indicated compound was added, and image acquisition was immediately started, obtaining an image every 7 s for a 30-minute period. Movies were processed and assembled using FIJI/Image J. Channels were gamma-adjusted to enhance visualization.

- Movie S1 – DMSO
- Movie S2 – Cilengitide
- Movie S3 – MK429 racemate
- Movie S4 – TDI-4161
- Movie S5 – TDI-3761

#### Supplementary Materials and Methods

##### HEK-293 Cells Expressing αVβ3:

HEK-293 cells were transfected using lipofectamine 2000 (Invitrogen #11668030) with the cDNA for αV using the pEF1/V5-His A vector and the cDNA for β3 using the vector pcDNA3.1. Cells expressing αVβ3 were identified by their reaction with murine monoclonal antibodies (mAbs) LM609[1] and 7E3,[2] and stable cell lines were established by repetitive sorting using LM609. Cells were found to not express αIIbβ3 as judged by negative reaction with mAb 10E5.[3,4] HEK-293 cells expressing αVβ3 (HEK-293-αVβ3 cells) for assays were counted in an automated cell counter (ADVIA 120) and adjusted to values appropriate for each assay.

##### αVβ3 Cell Adhesion to Fibrinogen Assay:

Polystyrene 96-well microtiter plates (Costar, 3590) were pre-coated with 3.5 µg/ml of purified fibrinogen (Enzyme Research Laboratories) in 0.15 M NaCl, 0.01 M Tris/HCl, pH 7.4 for 1 hour at 37°C. The wells were then washed and incubated with HEPES-Buffered Modified Tyrode's solution (HBMT; 0.128 M NaCl, 10 mM HEPES, 12 mM $NaHCO_3$, 0.4 mM $NaH_2P0_4$, 2.7 mM KCl, 0.35% bovine serum albumin (Fisher), 0.1% glucose, pH 7.4) for 1 hour at room temperature or overnight at 4°C. Wells were washed with HBMT containing 1 mM $Mg^{2+}$ and 2 mM $Ca^{2+}$ and then 50 µl of HEK-293-αVβ3 cells that were pretreated with the compound to be tested for 20 minutes at room temperature was added to each well at a concentration of 3,000 cells/µl. After ~30 minutes the wells were washed with HBMT containing $Ca^{2+}$ and $Mg^{2+}$ three times and then the adherent cells were lysed and the acid phosphatase activity that was released was measured by: 1. adding phosphatase substrate (Sigma P4744) at 2 mg/ml in 0.1 M Na citrate, pH 5.6, 0.1% Triton X-100 for 1 hour at room temperature; 2. stopping the reaction by adding 50 µl of 2 M NaOH; and 3. analyzing the samples in a spectrophotometer at 405 nm. In each assay, 10 mM EDTA was used as a positive control and untreated cells were used as a negative control. The $IC_{50}$ was defined as the concentration of the test compound that reduced the adhesion of the HEK-αVβ3 cells by 50%, taking the results with untreated cells as 100% and the results in the presence of EDTA as 0%.

##### AP5 Binding Assay:

HEK-αVβ3 cells were harvested using 0.05% trypsin, 0.5 mM EDTA, washed with HBMT once, and resuspended in HBMT containing 1 mM $Mg^{2+}$ and 2 mM $Ca^{2+}$. The cells were counted and adjusted to $5 \times 10^5$ cells per sample. The compound to be tested and fluorescently labeled mAb AP5 (either Alexa488 or Alexa674; 10 µg/ml) were added and incubated for 30 minutes at 37°C. The cells were then washed and analyzed by flow cytometry (BD FACSCalibur). In each assay, cilengitide (1 µM) and 10 mM EDTA were included as positive controls and untreated cells were used as the negative control. The concentration of the test compound required to induce the expression of 50% of the AP5 expression induced by 1 µM cilengitide was calculated and defined as the $EC_{50}$. The $EC_{50}$ for cilengitide was determined based on the expression induced by EDTA. The AP5 expression induced by 1 µM cilengitide was approximately twice the value with 10 mM EDTA [average ± SD of 17 experiments; control 7.6 ± 2.2, EDTA 21.8 ± 5.9, cilengitide 42.5 ± 8.0 arbitrary fluorescence units (AFU)]. In cases where even the highest concentration of test compound (10 µM) did not induce 50% exposure of the AP5 epitope, the results are reported as >10 µM.

**Priming Assay:**

HEK-293-αVβ3 cells were washed, resuspended in HBMT containing 1 mM $Mg^{2+}$ and 2 mM $Ca^{2+}$ at 2 x $10^6$/ml, and either left untreated (control) or incubated with 1 µM cilengitide, 100 µM RGDS, or 10 µM TDI-4161 or TDI-3761 for 20 minutes at room temperature. Samples were then fixed with an equal volume of 4% paraformaldehyde in PBS for 40 minutes at room temperature, followed by quenching of the reaction with twice the volume of 5 mM glycine for 5 minutes at room temperature. After washing 5x with HBMT, the cells were resuspended in HBMT containing 1 mM $Mg^{2+}$ and 2 mM $Ca^{2+}$. Alexa488-conjugated fibrinogen was then added (200 µg/ml final concentration) and incubated for 30 minutes at 37 °C. The cells were then washed once and analyzed by flow cytometry.

**De-adhesion Assays:**

We employed a modification of the assay reported by Charo et al. for human endothelial cells.[5] Adhesion of HEK-293-αVβ3 cells to fibrinogen was carried out as above for 30 minutes in the absence of compounds and unattached cells were removed by washing. Compounds were then added and after an additional 30-60 minutes, the microtiter wells were washed again and the number of remaining cells was analyzed and compared to the number of cells that adhered during the first 30 minutes. In some experiments, AP5 binding was performed on cells that de-adhered during the experiment in the presence of compound. Since cells in the control sample did not de-adhere during the additional 30-60 minutes, they were lifted from the plate by treatment with trypsin, but without EDTA. For time-lapse studies of the de-adhesion process, cells were plated on IbiTreat µ-Slide 8 well (ibidi, Martinsried, GmbH) pre-coated with fibrinogen as described above. Differential interference contrast (DIC) images were acquired on a Leica SP8 inverted confocal microscope at 12 bits in 1024 pixels × 1024 pixels using a water immersion objective Apo CS2 40x/1.10 numerical aperture, at 1 Airy unit pinhole diameter with manual laser-power intensity-compensation. For pre-treatment, images were taken every 7 s for a 5-minute period. After this time, the indicated compound was added, and image acquisition was immediately started obtaining an image every 7 s for a 30-minute period. Final images were processed and assembled using FIJI/Image J. For images and movies, channels were gamma-adjusted to enhance visualization.

**Mouse Endothelial Cell Adhesion Assay:**

To assess the reactivity of the αVβ3 antagonists with murine αVβ3, we tested the ability of the compounds to inhibit the adhesion of murine endothelial cells to immobilized fibrinogen. Mouse primary aortic endothelial cells were obtained from Cell Biologics and grown in endothelial cell medium (M1168). Cells were harvested with trypsin-EDTA, washed once with HBMT, and resuspended in HBMT containing 1 mM $Mg^{2+}$ and 2 mM $Ca^{2+}$. The cells were then incubated with the test compounds for 20 minutes at 22°C and added to microtiter wells that had been pre-coated by adding 50 µl of human fibrinogen (5 µg/ml) in Tris-saline buffer, pH 7.4 at 4°C overnight and then washing and blocking with HBMT containing 0.35% albumin. After incubating the cells in the wells for 30 minutes at 37°C, the wells were washed with HBMT containing $Mg^{2+}$ and $Ca^{2+}$ and the number of remaining adherent cells was assessed by lysing the cells in Triton X-100 and determining the acid phosphatase activity (Sigma P4744).

**Inhibition of Ligand Binding by Purified αVβ3 and αVβ5:**

The ability of compounds to interfere with the interaction of purified αVβ3 and αVβ5 with their ligands was tested by modifications of the assays described by Henderson et al.[6] The purified receptors were obtained from R&D Systems. The ligands employed were adenovirus 2 penton base[7] (kindly supplied by Dr. Glen Nemerow of Scripps Research Institute) for αVβ3 and vitronectin (Molecular Innovations # HVN) for αVβ5. Microtiter wells were coated overnight at 4°C with 50 µl of penton base or LAP at 0.2 µg/ml or vitronectin at 10 µg/ml, and then washed with TTBS buffer (137 mM NaCl, 25 mM Tris/HCl, 2.7 mM KCl, 0.1% Tween-20, pH 7.4) and blocked with 1% BSA for 60 minutes at 22°C. The purified receptor in TTBS + 0.1% BSA was incubated with the compound to be tested for 20 minutes at 22°C and then the mixture was added to the well. After 120 minutes at 37°C, the wells were washed with TTBS and the bound receptor measured by adding a biotin-labeled mAb to αV (R&D #BAF1219) at 0.1 µg/ml and detecting the antibody by using horse radish peroxidase-labeled streptavidin (GE Healthcare #RPN1231V). After subtracting the value obtained in the presence of EDTA (30 mM) from each result, the percentage inhibition of binding and the $IC_{50}$ for each compound was determined as above.

**Inhibition of Ligand Binding by αIIbβ3:**

The ability of the compounds to inhibit the interaction between αIIbβ3 and fibrinogen was measured by assessing their ability to inhibit the binding of HEK-293 cells expressing αIIbβ3 to immobilized fibrinogen as reported previously.[8]

**Osteoclast Culture for Morphology, Bone Lacunae Formation, and Cross-linked Collagen Degradation Peptides:**

All studies were performed on coded samples with the experimenter not knowing the identity of the individual compounds. As previously described,[9] bone marrow was harvested from the femur and tibia of 8-10 week old male C57BL mice using the α modification of Minimal Essential Medium (α-MEM) and cultured in 150-mm plastic petri dishes in α-MEM supplemented with 10% fetal bovine serum and 10% conditioned medium from the CMG14-12 cell line[10] as a source of M-CSF for 4 days. The resulting macrophages were collected with trypsin-EDTA and cultured in: plastic tissue culture plates with or without bovine bone slices (1.2 x $10^4$ cells/well/0.5 ml medium) with RANK ligand (100 ng/ml), M-CSF (2% CMG conditioned medium), test compounds, or DMSO (1:1000 final dilution). In some experiments, compounds were added on day 4 instead of on day 0. The medium was changed every two days and fresh aliquots of the compounds were added to the replacement medium. On day 5, cells grown in the plastic plate with bone were fixed with 4% paraformaldehyde for 10 min at room temperature and stained for tartrate-resistance acid phosphatase (Sigma). On day 6, the medium of cells grown on bone was collected for assay of cross-linked degradation products of collagen type 1 telopeptides (Crosslaps, IDS). After washing twice with PBS, the bone slices were transferred to a glass slide, and cover-slipped using 75% glycerol in water as a mounting medium. The specimens were visualized with a Nikon Eclipse-E400 fluorescent microscope. The cells grown on bone for osteoclast resorption lacunae analysis were removed from the bone slices on day 6 with mechanical agitation and the bone slices were incubated with peroxidase-conjugated wheat germ agglutinin (20 µg/ml) for 1 hour and then stained with 3,3'-diamaninobenzidine. Five defined 10x fields from the periphery (4) and center (1) were photographed and the percentage of the image containing lacunae was determined by image analysis (ImageJ). Each compound was studied on 5 separate bone slices.

In other experiments, murine bone marrow macrophages prepared as above were cultured with RANK ligand (100 ng/ml) and M-CSF in the presence of DMSO or test compounds (1:1000) for 3 days and then cells were lysed and tested for osteoclast differentiation markers (integrin subunit β3, NFATc1, and cathepsin K) by immunoblotting equal amounts of protein (30 μg). Bone marrow macrophages cultured in M-CSF alone served as a negative control.

**Mouse Aortic Ring Vascular Sprout Assay:**

These studies were conducted in accord with the policies of the Ethics Committee of Queen Mary University of London. 8 C57Bl6 mice were euthanized at 8-10 weeks of age by cervical dislocation and their thoracic aortae dissected. As previously described,[11] aortae were cut into 0.5 mm rings, starved overnight in OptiMEM medium with penicillin-streptomycin (GIBCO), and then embedded in 50 µl of 1.1 mg/ml rat tail collagen type 1 (Millipore). Collagen matrix was polymerized for 15 minutes at room temperature and subsequently for 1 hour at 37ºC. Aortic rings were then stimulated with 150 µl of OptimMEM supplemented with 2.5% FCS and 30 ng/ml VEGF, PBS, or vehicle (DMSO) alone. Cilengitide, TDI-4161, or TDI-3761 was added to OptimMEM supplemented with 2.5% FCS and 30 ng/ml VEGF at 1 nM, 10 nM, 100 nM, 1 μM and 10 μM. The number of sprouts per ring was counted using a phase contrast microscope at day 8 post embedding. After that, rings were fixed with 4% formaldehyde, permeabilized using 0.2% Triton X-100, blocked using 2% BSA, and stained with TRITC-conjugated lectin from Bandeiraea simplicifolia (BS-I; Sigma L9381/L5264). Images were captured using a LSM710 confocal microscope.

**Organic Synthesis of Select Compounds:**

**General features:**

Solvents and reagents were purchased from VWR or Sigma Aldrich. All reactions involving air- or moisture-sensitive compounds were performed under nitrogen atmosphere using dried glassware. $^{1}$H and $^{13}$C NMR spectra were recorded at 500 MHz and 125 MHz respectively, on a Bruker Advance III HD 500 MHz NMR spectrometer equipped with a TCI cryogenic probe with enhanced $^{1}$H and $^{13}$C detection. All data were collected at 298 °K and signals were reported in parts per million (ppm), internally referenced for $^{1}$H and $^{13}$C to chloroform signals at 7.26 ppm or 77.0 ppm, to DMSO signals at 2.50 ppm or 39.5 ppm, or TMS at 0 ppm. Chemical shifts are reported in ppm and the coupling constants ($J$) are expressed in hertz (Hz). Splitting patterns are designated as follows: s, singlet; d, doublet; t, triplet; m, multiplet; dd, doublet of doublets; ddd, doublet of doublets of doublets; and dt, doublet of triplets. Flash chromatography purifications were performed on Combi*Flash* Rf (Teledyne ISCO) as the stationary phase. Melting points were determined on an MP50 Melting Point System (Mettler Toledo). Purity for all tested compounds was determined through high-performance liquid chromatography and all compounds were found to be > 95% pure.

**Synthesis of individual compounds:**

**TDI-1366**

1. **Benzyl 3-amino-3-(6-methoxy-3-pyridyl)propanoate:** A mixture of 3-amino-3-(6-methoxy-3-pyridyl)propanoic acid (1.90 g, 9.68 mmol) and 4-methylbenzenesulfonic acid hydrate (2.03 g, 10.7 mmol) in phenylmethanol (10.47 g, 96.8 mmol) and toluene (30 mL) was stirred at 110 °C for 2 hours and then at 90°C overnight. The resulting clear solution was poured

into $NaHCO_3$ aq. and extracted with EtOAc. The organic layer was washed with brine and concentrated in vacuo. The residue was purified by silica gel column chromatography (0% - 10% MeOH in EtOAc) to give benzyl 3-amino-3-(6-methoxy-3-pyridyl)propanoate (1.06 g, 3.70 mmol, 38% yield) as a colorless oil. MS m/z: 288.1 $[M+H]^+$. $^1$H NMR (500 MHz, Chloroform-*d*) δ 8.14 (d, *J* = 2.5 Hz, 1H), 7.62 (dd, *J* = 8.6, 2.5 Hz, 1H), 7.41 – 7.31 (m, 5H), 6.74 (d, *J* = 8.5 Hz, 1H), 5.15 (s, 2H), 4.45 (dd, *J* = 8.6, 5.1 Hz, 1H), 3.95 (s, 3H), 2.80 – 2.65 (m, 2H). 2H not found.

2. **Benzyl 3-(6-methoxy-3-pyridyl)-3-[5-(5,6,7,8-tetrahydro-1,8-naphthyridin-2-yl)pentanoylamino]propanoate:** To a solution of 5-(5,6,7,8-tetrahydro-1,8-naphthyridin-2-yl)pentanoic acid (317.89 mg, 1.36 mmol), benzyl 3-amino-3-(6-methoxy-3-pyridyl)propanoate (370.00 mg, 1.29 mmol), HOBt (240.08 mg, 1.42 mmol) and triethylamine (326.90 mg, 3.23 mmol) in DMF (15 ml) was added 3-(ethyliminomethyleneamino)-N,N-dimethyl-propan-1-amine;hydrochloride (272.49 mg, 1.42 mmol) at room temperature. The mixture was stirred at room temperature overnight. The mixture was diluted with water and extracted with EtOAc. The organic layer was washed with $NaHCO_3$ aq. and brine, dried over $MgSO_4$ and concentrated in vacuo. The residue was purified by silica gel column chromatography (0-10% MeOH in EtOAc) to give benzyl 3-(6-methoxy-3-pyridyl)-3-[5-(5,6,7,8-tetrahydro-1,8-naphthyridin-2-yl)pentanoylamino]propanoate (600 mg, 1.19 mmol, 92% yield) as a white solid. $^1$H NMR (500 MHz, Chloroform-*d*) δ 8.11 (d, *J* = 2.5 Hz, 1H), 7.49 (dd, *J* = 8.6, 2.6 Hz, 1H), 7.35 (dd, *J* = 5.2, 1.9 Hz, 3H), 7.28 – 7.24 (m, 2H), 7.06 (d, *J* = 7.3 Hz, 1H), 6.68 (d, *J* = 8.6 Hz, 1H), 6.55 (d, *J* = 8.4 Hz, 1H), 6.35 (d, *J* = 7.3 Hz, 1H), 5.42 (dt, *J* = 8.4, 5.9 Hz, 1H), 5.13 – 5.02 (m, 2H), 4.80 (s, 1H), 3.94 (s, 3H), 3.40 (td, *J* = 5.7, 2.5 Hz, 2H), 3.02 – 2.94 (m, 2H), 2.93 – 2.82 (m, 2H), 2.70 (t, *J* = 6.4 Hz, 2H), 2.60 – 2.50 (m, 2H), 2.27 – 2.18 (m, 2H), 1.98 – 1.86 (m, 2H), 1.74 – 1.59 (m, 2H).

**3. 3-(6-Methoxy-3-pyridyl)-3-[5-(5,6,7,8-tetrahydro-1,8-naphthyridin-2-yl)pentanoylamino]propanoic acid (TDI-1366):** The mixture of benzyl 3-(6-methoxy-3-pyridyl)-3-[5-(5,6,7,8-tetrahydro-1,8-naphthyridin-2-yl)pentanoylamino]propanoate (95.00 mg, 189.02 µmol) and palladium (200.00 mg, 1.88 µmol) in MeOH (10 ml) was hydrogenated at room temperature overnight. The mixture was filtered through a Celite pad to remove Pd-C. The filtrate was concentrated in vacuo. The residue (diluted with CH3CN:water:MeOH=10:10:1) was purified by reversed phase chromatography (water:CH3CN= 0 - 60%) to give TDI-1366 (58.00 mg, 140.61 µmol, 74% yield) as a white solid. MS m/z: 413.4 $[M+H]^+$. $^1$H NMR (500 MHz, DMSO-$d_6$) δ 8.46 (s, 1H), 8.08 (d, *J* = 2.5 Hz, 1H), 7.65 (dd, *J* = 8.6, 2.5 Hz, 1H), 7.05 (d, *J* = 7.3 Hz, 1H), 6.77 (d, *J* = 8.5 Hz, 1H), 6.46 (s, 1H), 6.24 (d, *J* = 7.3 Hz, 1H), 5.11 (d, *J* = 6.8 Hz, 1H), 3.83 (s, 3H), 3.34 (s, 5H), 2.68 – 2.58 (m, 3H), 2.45 – 2.36 (m, 3H), 2.09 (td, *J* = 7.0, 2.9 Hz, 2H), 1.77 (p, *J* = 6.1 Hz, 2H), 1.50 (dt, *J* = 11.8, 5.3 Hz, 2H).

**TDI-2668**

1. **Methyl 4-[5-(5,6,7,8-tetrahydro-1,8-naphthyridin-2-yl)pentanoylamino]butanoate:**To a mixture of 5-(5,6,7,8-tetrahydro-1,8-naphthyridin-2-yl)pentanoic acid (50.00 mg, 213.41 µmol), methyl 4-aminobutanoate hydrochloride (49.17 mg, 320.12 µmol), HOBt H2O (49.02 mg, 320.12 µmol) and triethylamine (86.38 mg, 853.64 µmol, 118.33 µL) in DMF (2.0 ml) was added 3-(ethyliminomethyleneamino)-N,N-dimethyl-propan-1-amine;hydrochloride (61.37 mg, 320.12 µmol) at room temperature. The mixture was stirred at room temperature overnight. The mixture was concentrated in vacuo and the residue was purified

by NH-silica-gel column chromatography (30-100% AcOEt in hexane) to give methyl 4-[5-(5,6,7,8-tetrahydro-1,8-naphthyridin-2-yl)pentanoylamino]butanoate (71 mg). MS m/z: 334.2 $[M+H]^+$. $^1H$ NMR (500 MHz, Chloroform-*d*) δ 7.07 (d, *J* = 7.3 Hz, 1H), 6.36 (d, *J* = 7.3 Hz, 1H), 5.84 (t, *J* = 6.0 Hz, 1H), 4.85 (s, 1H), 3.69 (s, 3H), 3.42 (td, *J* = 5.7, 2.5 Hz, 2H), 3.30 (q, *J* = 6.6 Hz, 2H), 2.71 (t, *J* = 6.4 Hz, 2H), 2.60 – 2.53 (m, 2H), 2.39 (t, *J* = 7.2 Hz, 2H), 2.21 (d, *J* = 7.2 Hz, 2H), 1.96 – 1.89 (m, 2H), 1.85 (p, *J* = 7.1 Hz, 2H), 1.70 (dd, *J* = 7.7, 3.9 Hz, 4H).

2. **4-[5-(5,6,7,8-tetrahydro-1,8-naphthyridin-2-yl)pentanoylamino]butanoic acid (TDI-2668):** To a solution of methyl 4-[5-(5,6,7,8-tetrahydro-1,8-naphthyridin-2-yl)pentanoylamino]butanoate (71 mg) in MeOH (5.0 ml) was added NaOH (1M aq.) (5.00 mL) at room temperature. The mixture was stirred at room temperature overnight. The reaction was neutralized with 1M HCl and the mixture was concentrated in vacuo. The residue was purified by silica gel column chromatography (0-100% MeOH in $CH_2Cl_2$) and recrystallized from MeOH/AcOEt to give 4-[5-(5,6,7,8-tetrahydro-1,8-naphthyridin-2-yl)pentanoylamino]butanoic acid as a white solid (50 mg). MS m/z: 320.2 $[M+H]^+$. $^1H$ NMR (500 MHz, DMSO-*d*$_6$) δ 12.15 (brs, 1H), 7.81(s, 1H), 7.08(d, *J* = 6.9 Hz, 1H), 6.36(s, 1H), 6.28 (d, *J* = 6.9 Hz 1H), 3.31-3.22 (m, 2H), 3.09-3.01 (m, 2H), 2.63 (t, *J* = 4.3 Hz, 2H), 2.45 (t, *J* = 6.6 Hz, 2H), 2.23 (t, *J* = 7.1 Hz, 2H), 2.08 (t, *J* = 7.1 Hz, 2H), 1.82-1.74 (m, 2H), 1.67-1.45 (m, 6H).

**TDI-3761**

1. **Methyl 2-(benzyloxycarbonylamino)-4-[5-(5,6,7,8-tetrahydro-1,8-naphthyridin-2-yl)pentanoylamino]butanoate:** To a solution of 5-(5,6,7,8-tetrahydro-1,8-naphthyridin-2-yl)pentanoic acid (2.80 g, 11.95 mmol), 3-(ethyliminomethyleneamino)-N,N-dimethyl-propan-1-amine hydrochloride (4.58 g, 23.90 mmol), HOBt dihydrate (4.09 g, 23.90 mmol) and DIPEA (6.18 g, 47.80 mmol, 8.35 ml) in DMF (50 ml) was added methyl 4-amino-2-(benzyloxycarbonylamino)butanoate hydrochloride (3.62 g, 11.95 mmol) at room temperature. The mixture was stirred at room temperature overnight, then quenched with water, diluted with AcOEt, washed with water and brine, dried with $MgSO_4$ and concentrated in vacuo. The residue was purified by chromatography silica gel (hexane:AcOEt = 10–100%, AcOEt:MeOH = 0–20%) to give methyl 2-(benzyloxycarbonylamino)-4-[5-(5,6,7,8-tetrahydro-1,8-naphthyridin-2-yl)pentanoylamino]butanoate (4.14 g, 8.58 mmol, 71% yield) as a white solid. MS m/z: 483.3 $[M+H]^+$. $^1H$ NMR (500 MHz, DMSO-*d*$_6$) δ 7.84 (t, *J* = 5.5 Hz, 1H), 7.77 (d, *J* = 7.8 Hz, 1H), 7.47 – 7.23 (m, 5H), 7.02 (d, *J* = 7.2 Hz, 1H), 6.31 – 6.16 (m, 2H), 5.11 – 4.99 (m, 2H), 4.08 (ddd, *J* = 9.7, 7.8, 4.8 Hz, 1H), 3.63 (s, 3H), 3.24 (dq, *J* = 6.0, 2.8 Hz, 2H), 3.15 – 3.01 (m, 2H), 2.61 (t, *J* = 6.3 Hz, 2H), 2.46 – 2.35 (m, 2H), 2.06 (t, *J* = 7.1 Hz, 2H), 1.93 – 1.65 (m, 4H), 1.51 (dq, *J* = 22.3, 7.6 Hz, 4H).

2. **Methyl 2-amino-4-[5-(5,6,7,8-tetrahydro-1,8-naphthyridin-2-yl)pentanoylamino]butanoate:** The mixture of methyl 2-(benzyloxycarbonylamino)-4-[5-(5,6,7,8-tetrahydro-1,8-naphthyridin-2-yl)pentanoylamino]butanoate (2.43 g, 5.04 mmol) and Pd/C (250.00 mg, 2.06 mmol) in MeOH (25 ml) was stirred under $H_2$ atmosphere (1 atm) at room temperature overnight. The mixture was filtrated (celite) and concentrated in vacuo. To the residue, AcOEt and hexanes were added and the crystal was collected by filtration to give methyl 2-amino-4-[5-(5,6,7,8-tetrahydro-1,8-naphthyridin-2-yl)pentanoylamino]butanoate (1.90 g). MS m/z: 449 $[M+H]^+$. $^1H$ NMR (500 MHz, DMSO-*d*$_6$) δ 7.80 (t, *J* = 5.6 Hz, 1H), 7.03 (d, *J* = 7.2 Hz, 1H), 6.32 – 6.21 (m, 2H), 3.62 (s, 3H), 3.35 (s, 3H), 3.25 (dq, *J* = 6.0, 2.7 Hz, 2H), 3.13 (dt, *J* = 7.4, 6.2 Hz,

2H), 2.61 (t, *J* = 6.3 Hz, 2H), 2.42 (t, *J* = 7.3 Hz, 2H), 2.06 (t, *J* = 7.2 Hz, 2H), 1.87 – 1.70 (m, 4H), 1.60 – 1.42 (m, 4H).

3. **2-(1,3-Benzoxazol-2-ylamino)-4-[5-(5,6,7,8-tetrahydro-1,8-naphthyridin-2-yl)pentanoylamino]butanoic acid (TDI-3761):** To a solution of methyl 2-amino-4-[5-(5,6,7,8-tetrahydro-1,8-naphthyridin-2-yl)pentanoylamino]butanoate (101.20 mg, 290.44 µmol) and triethylamine (58.78 mg, 580.88 µmol, 80.52 µl) in THF (6.0 ml) was added 2-chlorobenzoxazole (89.21 mg, 580.88 µmol, 66.32 µl) at room temperature. The mixture was stirred at room temperature for 1 hour, then warmed to 60 °C and stirred overnight. The mixture was then stirred at 80 °C for 2 days. The reaction mixture was filtered and concentrated in vacuo. The residue was purified by NH silica gel chromatography (hexane:AcOEt = 80:20 to 0:100) to give methyl 2-(1,3-benzoxazol-2-ylamino)-4-[5-(5,6,7,8-tetrahydro-1,8-naphthyridin-2-yl)pentanoylamino]butanoate (62.40 mg, 134.04 µmol, 46% yield) as a colorless oil. To a solution of methyl 2-(1,3-benzoxazol-2-ylamino)-4-[5-(5,6,7,8-tetrahydro-1,8-naphthyridin-2-yl)pentanoylamino]butanoate (62.40 mg, 134.04 µmol) in THF (4.00 ml) and water (1.00 mL) was added LiOH ($H_2O$) (16.87 mg, 402.12 µmol) at 0°C. The mixture was stirred at room temperature for 4 days. The reaction mixture was quenched with 1N HCl at 0°C and concentrated in vacuo. The residue was purified by preparative thin layer chromatography (PTLC) (EtOAc:MeOH = 1:1) and the silica gel was extracted with MeOH and concentrated in vacuo to give 2-(1,3-benzoxazol-2-ylamino)-4-[5-(5,6,7,8-tetrahydro-1,8-naphthyridin-2-yl)pentanoylamino]butanoic acid (57.10 mg, 126.46 µmol, 94% yield) as an amorphous solid. MS m/z: 452 $[M+H]^+$. 1H NMR (500 MHz, Methanol-d4) δ 7.22 – 7.12 (m, 2H), 7.11 – 7.02 (m, 2H), 6.94 (dd, J = 7.8, 1.3 Hz, 1H), 6.29 (d, J = 7.3 Hz, 1H), 4.18 (dd, J = 7.8, 4.7 Hz, 1H), 3.29 (dd, J = 4.7, 2.9 Hz, 2H), 3.24 (m, 7H), 2.61 (t, J = 6.4 Hz, 2H), 2.46 (d, J = 7.1 Hz, 2H), 2.18 – 2.06 (m, 3H), 1.95 – 1.88 (m, 1H), 1.80 – 1.73 (m, 2H), 1.56 (dt, J = 7.1, 3.6 Hz, 4H). 1H not found.

**TDI-3909**

1. **2-Amino-4-[5-(5,6,7,8-tetrahydro-1,8-naphthyridin-2-yl)pentanoylamino]butanoic acid:** To a mixture of methyl 4-[5-(5,6,7,8-tetrahydro-1,8-naphthyridin-2-yl)pentanoylamino]butanoate (100.00 mg, 286.99 µmol), $H_2O$ (500 µl), and THF (2.0 ml) was added lithium hydroxide monohydrate (36.13 mg, 860.98 µmol) at room temperature. After being stirred at room temperature overnight, 1N HCl (1.0 M, 860 µl) was added to the reaction mixture. The mixture was concentrated in vacuo to give 2-amino-4-[5-(5,6,7,8-tetrahydro-1,8-naphthyridin-2-yl)pentanoylamino]butanoic acid as a white solid. This product was used in the next reaction without purification.

2. **2-(1,3-benzothiazole-2-carbonylamino)-4-[5-(5,6,7,8-tetrahydro-1,8-naphthyridin-2-yl)pentanoylamino]butanoic acid (TDI-3909):** To a suspension of 1,3-benzothiazole-2-carboxylic acid (394.24 mg, 2.20 mmol) in $CH_2Cl_2$ (10 ml) was added $(COCl)_2$ (418.87 mg, 3.30 mmol, 279.25 µl) and one drop of DMF on ice. The mixture was stirred at room temperature for 3 hours, then concentrated in vacuo. 5 mL of DMF was added to the residue to give an acid chloride solution. To a mixture of 2-amino-4-[5-(5,6,7,8-tetrahydro-1,8-naphthyridin-2-yl)pentanoylamino]butanoic acid (407.96 mg) in DMF (10 ml) were added the acid chloride solution and TEA (222.62 mg, 2.20 mmol, 304.96 µl) at room temperature. The mixture was stirred at room temperature for 2 hours. The reaction was quenched with water and saturated $NaHCO_3$ and then concentrated in vacuo. To the residue, MeOH and $CH_2Cl_2$ were added and the solution

filtered. The filtrate was concentrated in vacuo. The residue was purified by silica gel column chromatography (0-50% MeOH in $CH_2Cl_2$) followed by crystallization from MeOH/EtOAc to give 2-(1,3-benzothiazole-2-carbonylamino)-4-[5-(5,6,7,8-tetrahydro-1,8-naphthyridin-2-yl)pentanoylamino]butanoic acid (74.60 mg, 150.53 µmol, 14% yield) as a white solid. MS m/z: 496 $[M+H]^+$. $^1$H NMR (500 MHz, DMSO-$d_6$) δ 9.16 (d, *J* = 7.7 Hz, 1H), 8.26 (d, *J* = 8.0 Hz, 1H), 8.19 (d, *J* = 8.0 Hz, 1H), 7.88 (t, *J* = 5.5 Hz, 1H), 7.70 – 7.58 (m, 2H), 7.11 (d, *J* = 7.2 Hz, 1H), 6.95 (brs, 1H), 6.30 (d, *J* = 7.2 Hz, 1H), 4.45-4.37 (m, 1H), 3.27 (t, *J* = 5.6 Hz, 2H), 3.24-3.08 (m, 2H), 2.62 (t, *J* = 6.3 Hz, 2H), 2.45 (t, *J* = 7.4 Hz, 2H), 2.13 – 1.97 (m, 4H), 1.81 – 1.72 (m, 2H), 1.60-1.45 (m, 4H).

**TDI-4161**

1. **Methyl (*S*)-4-amino-2-{[(benzyloxy)carbonyl]amino}butanoate hydrochloride:** To a solution of hydrogen chloride in methanol (4 M, 140 ml, 10.0 eq) at 25°C was added compound (*S*)-4-amino-2-{[(benzyloxy)carbonyl]amino}butanoic acid (14.0 g, 55.50 mmol, 1.0 eq). Then the reaction mixture was stirred at 25°C for 10 hours. LC/MS showed all starting material was consumed and the desired product was detected. The reaction mixture was concentrated in *vacuo* to give the crude methyl (*S*)-4-amino-2-{[(benzyloxy)carbonyl]amino}butanoate hydrochloride (15.0 g) as a yellow oil that was used in the next step without further purification. MS m/z: 267.1 $[M+H]^+$. $^1$H NMR (400 MHz, $CD_3OD$) δ 7.3–7.31 (m, 5H), 5.11 (s, 2H), 4.34–4.25 (m, 2H), 3.89 (s, 2H), 3.74 (s, 1H), 3.31–3.20 (m, 1H), 3.03 (t, *J* = 7.6 Hz, 2H), 2.31–2.24 (m, 2H), 2.23–2.05 (m, 1H).

2. **Methyl (2S)-2-(benzyloxycarbonylamino)-4-[5-(5,6,7,8-tetrahydro-1,8-naphthyridin-2-yl)pentanoylamino]butanoate:** To the solution of 5-(5,6,7,8-tetrahydro-1,8-naphthyridin-2-yl)pentanoic acid (7.00 g, 29.88 mmol, 1.0 eq), HATU (22.72 g, 59.76 mmol, 2.0 eq) and N,N-diisopropylethylamine (DIPEA) (15.45 g, 119.52 mmol, 20.88 mL, 4.0 eq) in N,N-dimethylformamide (50 mL) was added compound (*S*)-4-amino-2-[(benzyloxy)carbonyl]amino}butanoate hydrochloride (14.21 g, 29.88 mmol, 1.0 eq) at 25°C. The mixture was stirred for 4 hours at 25°C. LC/MS showed the reaction was completed. The residue was poured into ice-water (150 ml). The mixture was extracted with ethyl acetate (100 mL x3). The combined organic phases were washed with brine (100 ml x3), dried over anhydrous sodium sulfate, filtered, and concentrated in vacuum to give compound methyl (2S)-2-(benzyloxycarbonylamino)-4-[5-(5,6,7,8-tetrahydro-1,8-naphthyridin-2-yl)pentanoylamino]butanoate (20.0 g) as yellow oil. MS m/z: 483.1 $[M+H]^+$. $^1$H NMR (400 MHz, $CDCl_3$) δ 7.40 (d, *J* = 7.6 Hz, 1H), 7.33–7.29 (m, 7H), 6.90 (br. s, 1H), 6.69 (br. s, 1H), 5.07 (s, 2H), 4.38–4.34 (m, 1H), 3.51–3.48 (m, 2H), 3.17–2.94 (m, 2H), 2.78 (s, 3H), 2.76–2.73 (m, 2H), 2.71–2.65 (m, 2H), 2.27–2.23 (m, 2H), 2.11–2.02 (m, 2H), 1.94–1.83 (m, 2H), 1.70–1.53 (m, 4H).

**3. Methyl (*S*)-2-amino-4-[5-(5,6,7,8-tetrahydro-1,8-naphthyridin-2-yl)pentanamido]butanoate:** To a solution of compound methyl (2S)-2-(benzyloxycarbonylamino)-4-[5-(5,6,7,8-tetrahydro-1,8-naphthyridin-2-yl)pentanoylamino]butanoate (20 g, 41.44 mmol, 1.0 eq) in methanol (200 ml) was added $Pd(OH)_2$ (1.16 g, 4.14 mmol, 50% purity, 0.1 eq) and the mixture was stirred for 6 hours under $H_2$ (15 psi) at 25°C. The reaction mixture was filtered and the filtrate was concentrated in vacuo to give the crude product methyl (*S*)-2-amino-4-[5-(5,6,7,8-tetrahydro-1,8-naphthyridin-2-

yl)pentanamido]butanoate (14.0 g) as a yellow oil that was used in the next step without further purification. MS m/z: 349.1 $[M+H]^+$.

4. **Methyl (*S*)-2-(benzo[*d*]thiazole-2-carboxamido)-4-(5-(5,6,7,8-tetrahydro-1,8-naphthyridin-2-yl)pentanamido)butanoate:** To a solution of compound methyl (*S*)-2-amino-4-[5-(5,6,7,8-tetrahydro-1,8-naphthyridin-2-yl)pentanamido]butanoate (6.00 g, 17.22 mmol, 1.0 eq) and compound benzo[*d*]thiazole-2-carboxylic acid (3.39 g, 18.94 mmol, 1.1 eq) in N,N-dimethylformamide (50 mL) was added DIPEA (6.68 g, 51.66 mmol, 9.03 ml, 3.0 eq) and [Bis(dimethylamino)methylene-1H-1,2,3-triazolo[4,5-b]pyridinium 3-oxid hexafluorophosphate (HATU) (13.09 g, 34.44 mmol, 2.0 eq) at 25°C. Then the mixture was stirred for 5 hours. The reaction mixture was poured into ice-water (150 ml). The aqueous phase was extracted with ethyl acetate (80 ml) x3. The combined organic phases were washed with brine (80 ml) 3, dried over anhydrous sodium sulfate, filtered, and concentrated in vacuo. The residue was purified by silica gel chromatography (dichloromethane: methanol = 200:1 to 10:1) to give compound methyl (*S*)-2-(benzo[*d*]thiazole-2-carboxamido)-4-(5-(5,6,7,8-tetrahydro-1,8-naphthyridin-2-yl)pentanamido)butanoate (5.00 g, 8.83 mmol, 51% yield) as a yellow oil. MS m/z: 510.2 $[M+H]^+$

**5. (2S)-2-(1,3-benzothiazole-2-carbonylamino)-4-[5-(5,6,7,8-tetrahydro-1,8-naphthyridin-2-yl)pentanoylamino]butanoate (TDI-4161):** Lithium hydroxide monohydrate (2.11 g, 88.30 mmol, 5.0 eq) was added into the solution of compound methyl (*S*)-2-(benzo[*d*]thiazole-2-carboxamido)-4-(5-(5,6,7,8-tetrahydro-1,8-naphthyridin-2-yl)pentanamido)butanoate (9.0 g, 17.66 mmol, 1.0 eq) in tetrahydrofuran (100 ml) and stirred for 1 hour at 20°C. TLC (dichloromethane: methanol= 10:1) showed the starting material was consumed completely and the desired product was detected. The reaction mixture was diluted with water (100 mL) and then washed with ethyl acetate (50 ml). The aqueous phase was acidified by 1N HCl solution to pH = 7 and the product was precipitated. The mixture was filtered and dried under vacuum to afford the crude product [(5.1 g; 53% yield; 90% purity; 93% enantiomeric excess (ee)]. The crude product (5.0 g) was dissolved in MeOH (1.0 L) and the suspension was heated to 80°C for 1 hour until the solid was dissolved completely. Then the mixture was cooled to 25°C gradually and stirred for 10 hours at this temperature. The white precipitated solid was filtered to give 3 g of desired product and the mother liquid was concentrated to half volume and some of product was precipitated out again. The solid was filtered and combined the previous batch (3 g) to afford the desired product (2S)-2-(1,3-benzothiazole-2-carbonylamino)-4-[5-(5,6,7,8-tetrahydro-1,8-naphthyridin-2-yl)pentanoylamino]butanoate (4.61 g, 9.27 mmol, 92% yield, 99.6% purity, 100% ee) as an off-white solid. MS m/z: 496.1 $[M+H]^+$. $^1H$ NMR (400 MHz, $CDCl_3$) δ 10.40 (br. s, 1H), 8.59 (d, *J* = 6.4 Hz, 1H), 8.12 (d, *J* = 8.0 Hz, 1H), 7.97 (d, *J* = 7.6 Hz, 2H), 7.54 (t, *J* = 7.2 Hz, 1H), 7.50-7.48 (t, *J* = 3.2 Hz, 1H), 7.28 (s, 1H), 6.32 (d, *J* = 7.2 Hz, 1H), 4.52-4.48 (m, 1H), 3.80–3.75 (m, 2H), 3.54 (t, *J* = 5.2 Hz, 2H), 3.41–3.35 (m, 1H), 2.86–2.79 (m, 1H), 2.75 (t, *J* = 6.0 Hz, 2H), 2.56–2.50 (m, 1H), 2.34–2.32 (m, 1H), 2.20–2.15 (m, 2H), 1.96–1.87 (m, 4H), 1.74–1.71 (m, 2H).

#### Supplementary Table

**Table S1. Data collection and refinement statistics.**

| | |
|---|---|
| PDB Code | **6MK0** |
| ***Data collection*** | **αVβ3-TDI4161** |
| Beamline | ID19 at APS |
| Space group | $P3_221$ |
| Unit cell dimensions (Å, º) | *a*=*b*=129.4, *c*=305.6;<br>α=β=90, γ=120 |
| Resolution range (Å) | 50-3.0 |
| Wavelength (Å) | 0.97895 |
| Total reflections | 1,632,693 |
| Unique reflections | 59,916 (5,883)* |
| Completeness | 100 (100) |
| Redundancy | 7.4 (6.3) |
| Molecules in asymmetric unit | 1 |
| Average $I/\sigma$ | 23.2 (2.4) |
| $R_{merge}$ (%) | 13.6 (163) |
| $R_{meas}$ (%) | 14.5 (136) |
| $R_{pim}$ (%) | 5.2 (53.7) |
| $CC_{1/2}$ | 0.98 (0.59) |
| Wilson *B*-factor | 49.4 |
| ***Refinement statistics*** | |
| Resolution range (Å) | 50-3.0 |
| $R_{factor}$ (%) | 22.5 (30.3) |
| $R_{free}$ (%)# | 25.5 (33.6) |
| No. of atoms | |
| Protein residues | 1,629 |
| Water | 4 |
| $Mn^{2+}$ | 8 |
| Glc-NAc | 14 |
| Average *B*-factor | |
| for all atoms ($Å^2$) | 54.6 |
| for the receptor | 54.4 |
| for TDI-4161 | 63.8 |
| for solvent ($Å^2$) | 38.6 |
| r.m.s. deviations | |
| Bond lengths (Å) | 0.003 |
| Bond angles (°) | 0.86 |
| Ramachandran plot | |
| Mos favored (%) | 89.0 |
| Allowed regions (%) | 10.6 |
| Outliers (%) | 0.4 |
| Clashscore (%) | 8.4 |
| Rotamer outliers (%) | 1.7 |

* Values in parentheses are for the highest resolution shell (0.1Å)
### $R_{free}$ was calculated with 5% of the data

#### Supplementary Figures

**Figure S1. Root Mean Square Deviation (RMSD) of MK-429 heavy atoms during three independent MD simulations of the ligand-receptor system.**

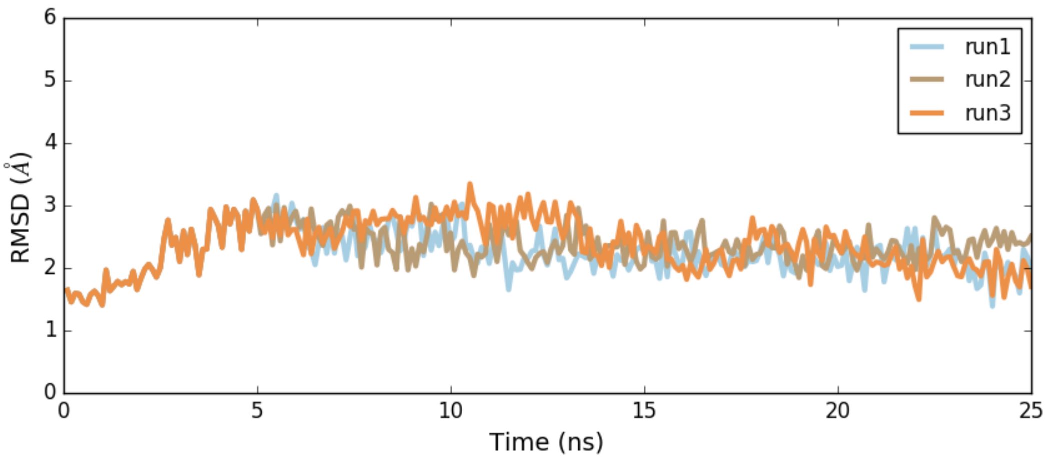

**Figure S2. RMSD of TDI-4161 heavy atoms during three independent MD simulations of the ligand-receptor system.**

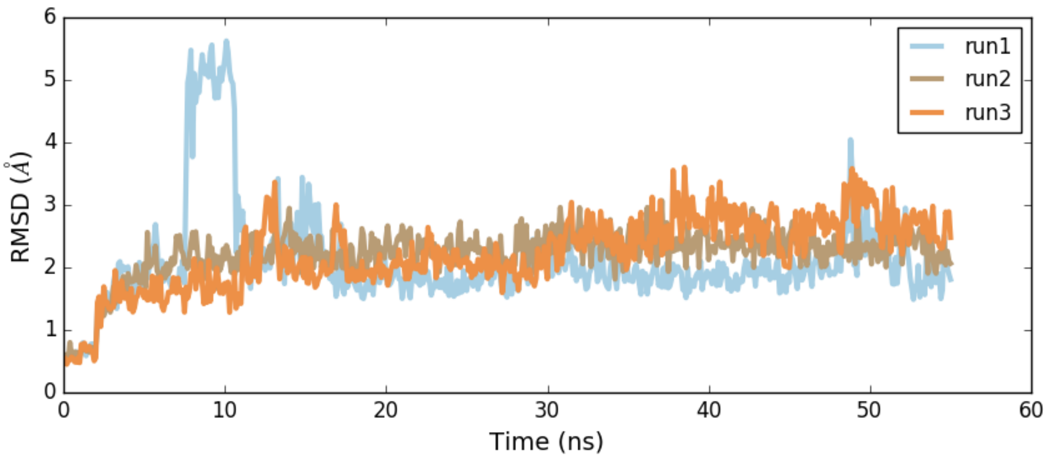

**Figure S3. Center of mass (COM) distances measured from three independent MD simulations of MK-429 bound to αVβ3**. **(A)** Distance between MK-0429 methoxypyridine moiety aromatic ring COM and β3 Y122 aromatic ring COM. **(B)** Distance between MK-429 tetrahydronaphthyridin moiety nitrogen atoms COM and αV D218 sidechain oxygen atoms COM. **(C)** Distance between MK-429 carboxyl terminal oxygen atoms COM and MIDAS ion.

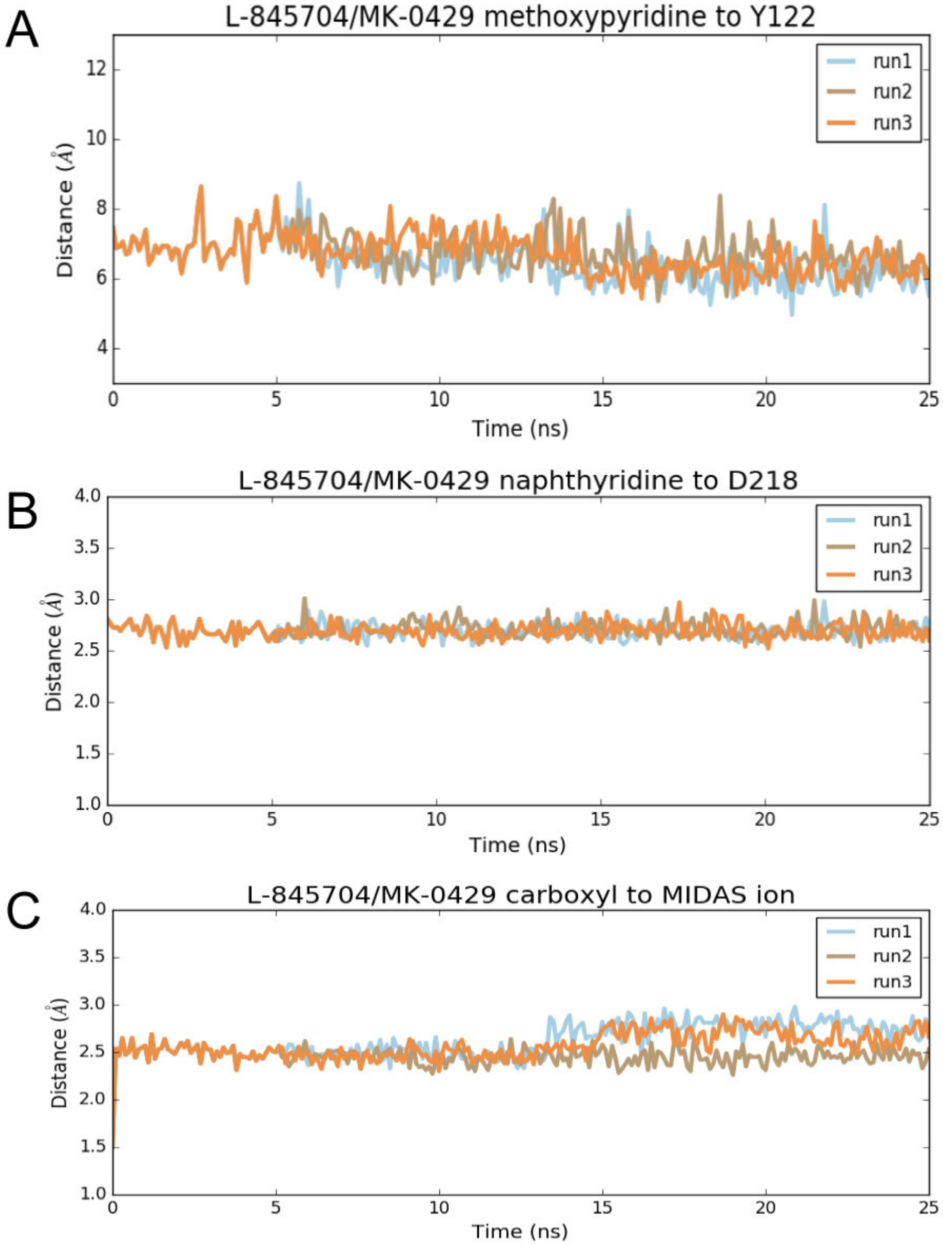

**Figure S4. Center of mass (COM) distances measured from three independent MD simulations of TDI-4161 bound αVβ3. (A)** Distance between TDI-4161 benzothiazole moiety COM and β3 Y122 aromatic ring COM. **(B)** Distance between TDI-4161 tetrahydronaphthyridin moiety nitrogen atoms COM and αV D218 sidechain oxygen atoms COM. **(C)** Distance between TDI-4161 carboxyl terminal oxygen atoms COM and MIDAS ion.

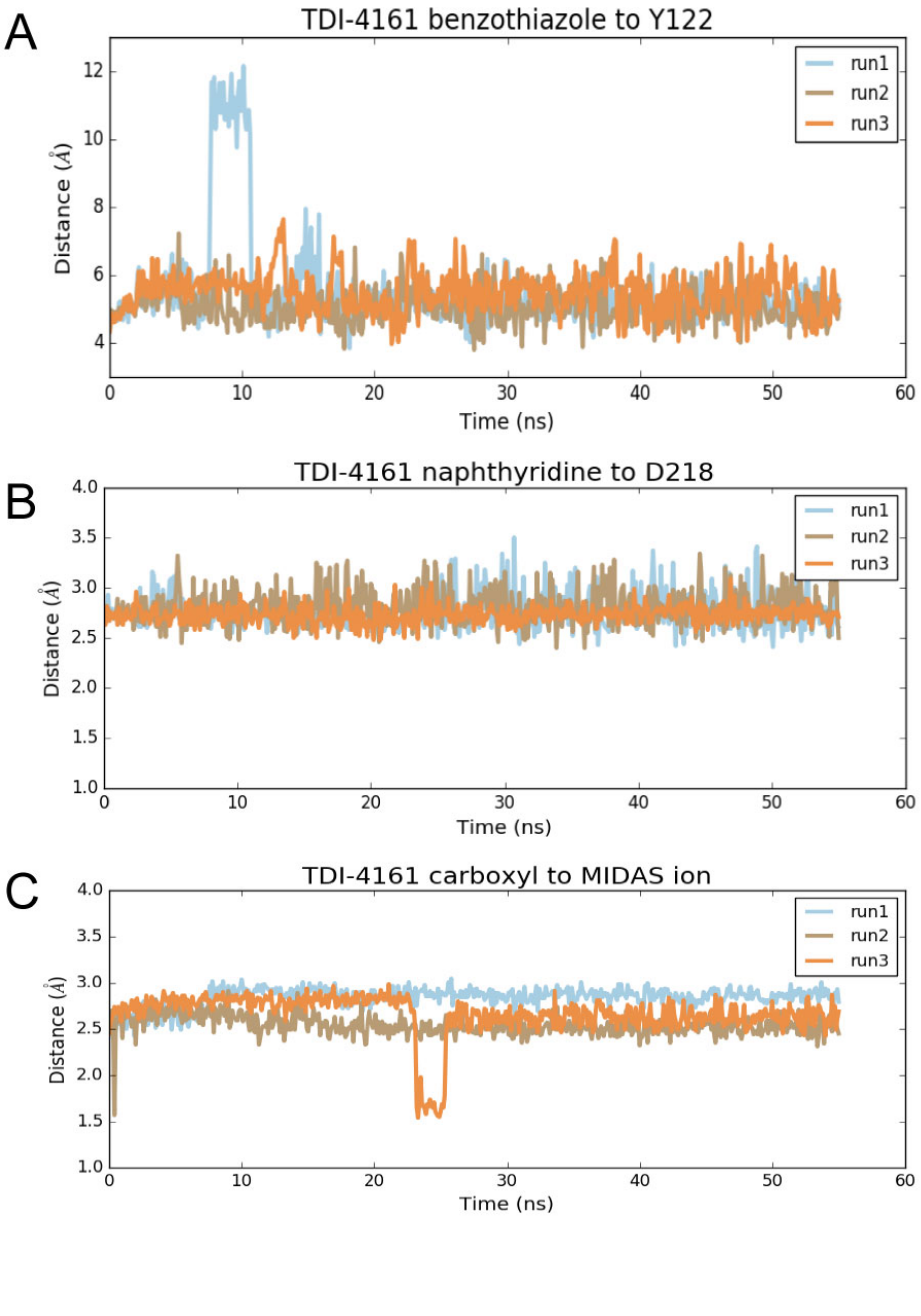

**Figure S5. Overlap between the predicted docking pose and crystal structure of TDI-4161 to αVβ3.** The predicted docking pose and crystal structure of TDI-4161 are shown in light grey and cyan colors, respectively. The αV and β3 backbones are shown in blue and red cartoon representations, respectively. Side chains of αV-D218 and β3-Y122 are shown as sticks. The MIDAS metal ion is shown as a purple sphere.

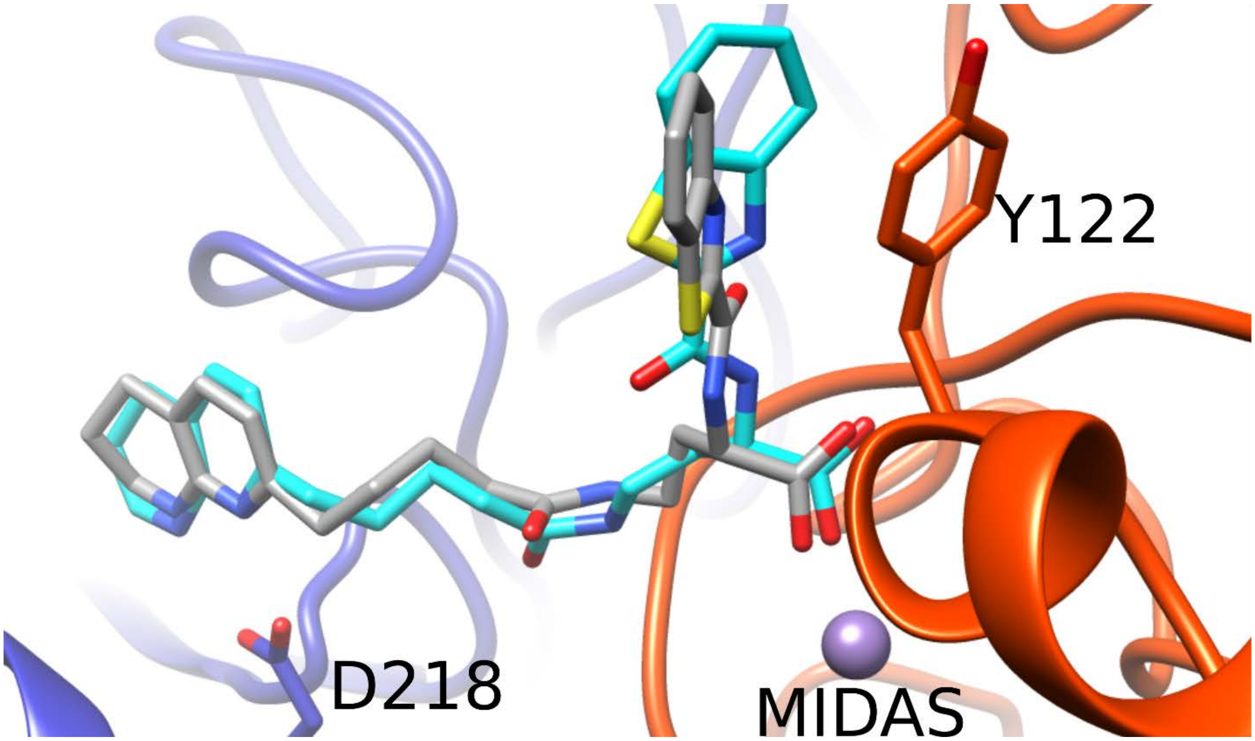

**Figure S6. The βA/βTD interface in the crystal structures of the αVβ3–TDI-4161 and αVβ3–wild-type (wt) FN10 complexes.** (A, B) 2Fo−Fc maps contoured at 1.0 σ for βTD residues 671-676 and Q319 of βA in the crystal structures of the αVβ3–TDI-4161 complex (A) and the αVβ3–wtFN10 complex (B). The propeller domain is in light blue and the βA domains in the αVβ3–TDI-4161 and αVβ3–wtFN10 complexes are in copper and rose, respectively. TDI-4161 and wtFN10 are shown as cyan and wheat sticks, respectively. Metal ions at LIMBS, MIDAS, and ADMIDAS are shown as grey, cyan, and magenta spheres, respectively. By blocking the inward movement of the α1 helix (by interacting with Y122), TDI-4161 stabilizes the contact between the carbonyl O of M335 of the F/α7 loop with the ADMIDAS metal ion, which is associated with a hydrogen bond between Nε2 of Q319 with Oγ of S674 of the βTD domain that allows for visualization of the glycan NAG711 at N654 of the βTD.

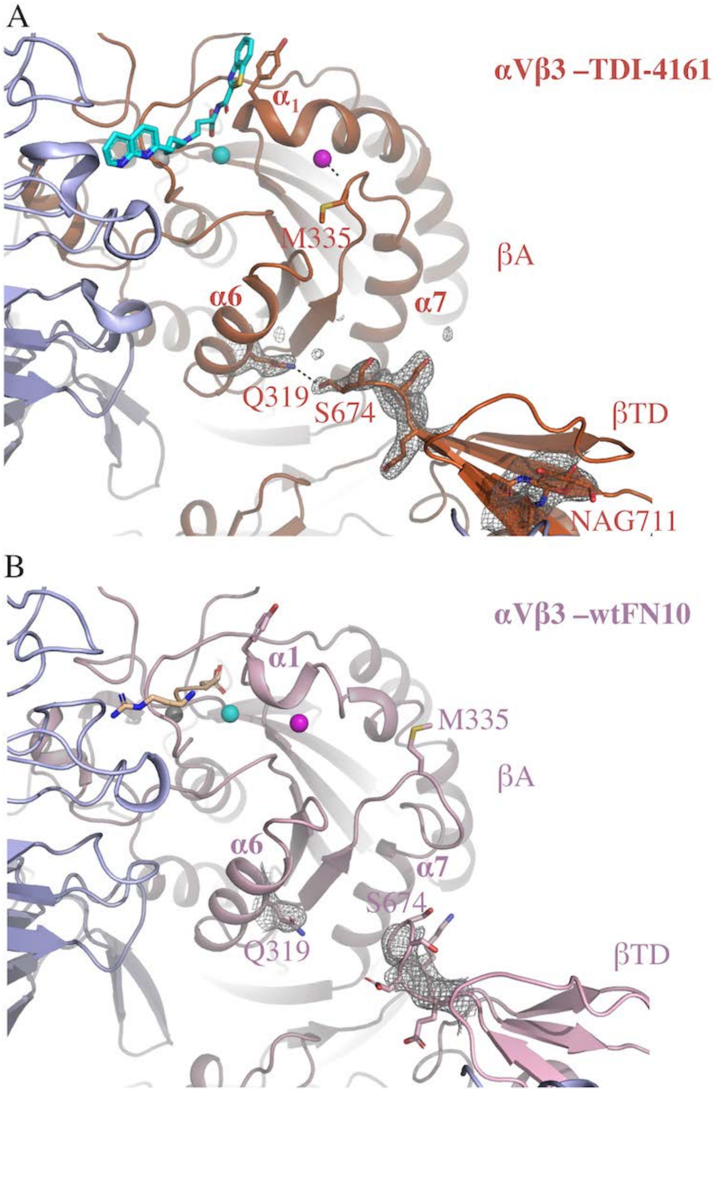

**Figure S7. Negative-stain EM analysis of the effect of αVβ3 antagonists on αVβ3 conformation.** (A-E) The 100 class averages of αVβ3 molecules in the presence of 0.1% DMSO (vehicle control) (A) or 10 µM of cilengitide (B), MK-429 (C), TDI-4161 (D), or TDI-3761 (E). Class averages in the compact-closed conformation are indicated by a red border; those in the extended-closed conformation by a blue border; and those in the extended-open conformation by green border. F. Number of particles analyzed (Total), number of particles that could not be unambiguously assigned (Unassigned), and number of particles in each of the three integrin conformations for each sample.

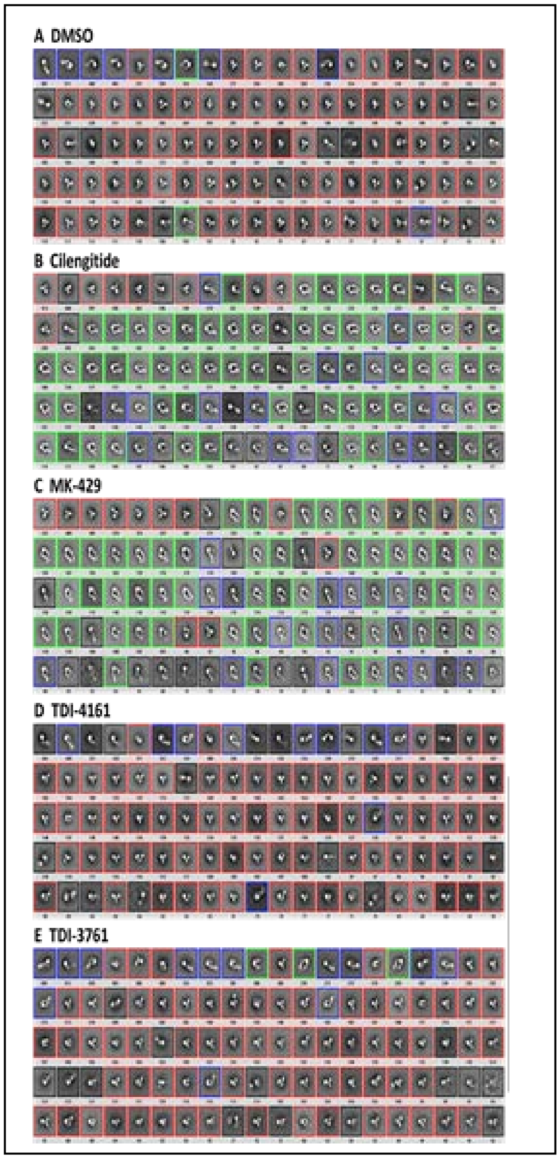

F.

| | DMSO | Cilengitide | MK-429 | TDI-4161 | TDI-3761 |
|---|---|---|---|---|---|
| Total | 18,693 | 17,345 | 14,933 | 15,373 | 17,302 |
| Unassigned | 1,946 | 2,358 | 1,760 | 3,045 | 1,522 |
| Compact Closed | 12,852 | 2,966 | 3,622 | 9,780 | 10,556 |
| Extended Closed | 3,438 | 2,022 | 1,702 | 2,548 | 4,377 |
| Extended Open | 457 | 9,999 | 7,849 | 0 | 847 |

**Figure S8. Effect of TDI-4161 and TDI-3761 on adhesion of HEK-293 cells expressing αIIbβ3 to immobilized fibrinogen.** At 10 μM neither compound inhibited αIIbβ3-mediated cell adhesion, nor did the αVβ3 antagonist cilengitide at 10 μM, the anti-αVβ3 mAb LM609 at 20 μg/ml, or the vehicle control (0.1% DMSO). In contrast, EDTA (10 mM), mAbs 10E5 and 7E3 (20 μg/ml of each), and the αIIbβ3 antagonist tirofiban (2 μM) all dramatically inhibited the adhesion. Data are expressed as a percentage of the adhesion of the untreated control cells and presented as mean ± SD of 3 separate experiments, each conducted in triplicate.

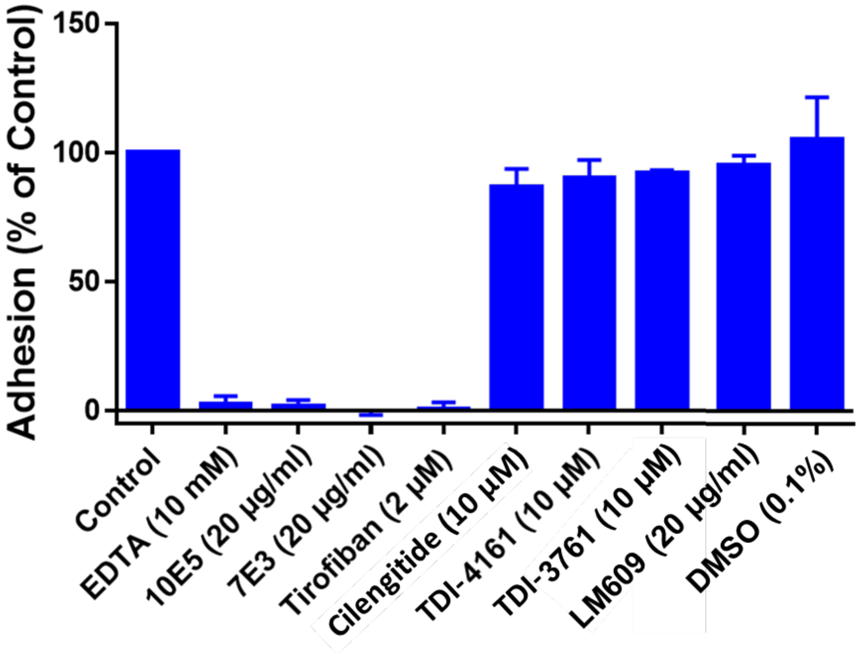

**Figure S9. De-adhesion of HEK-αVβ3 cells from fibrinogen.** (A) Adhesion of cells to fibrinogen when compounds were added either before the cells were added to the microtiter wells (blue bars) or 30 minutes after the cells were added (red bars). In the former case, the wells were evaluated 30 minutes after adding the cells. In the latter case, the wells were evaluated 30 minutes after the compounds were added to the wells containing the adherent cells (n = 3). (B) Exposure of the AP5 epitope on cells treated with compounds in suspension before adding cells to the microtiter wells (blue bars) or on cells that came off the well after adding compounds with washing (MK-429 racemate, TDI-4161, TDI-3761, and cilengitide) or by being treated with trypsin (control compound and DMSO) (n = 3).

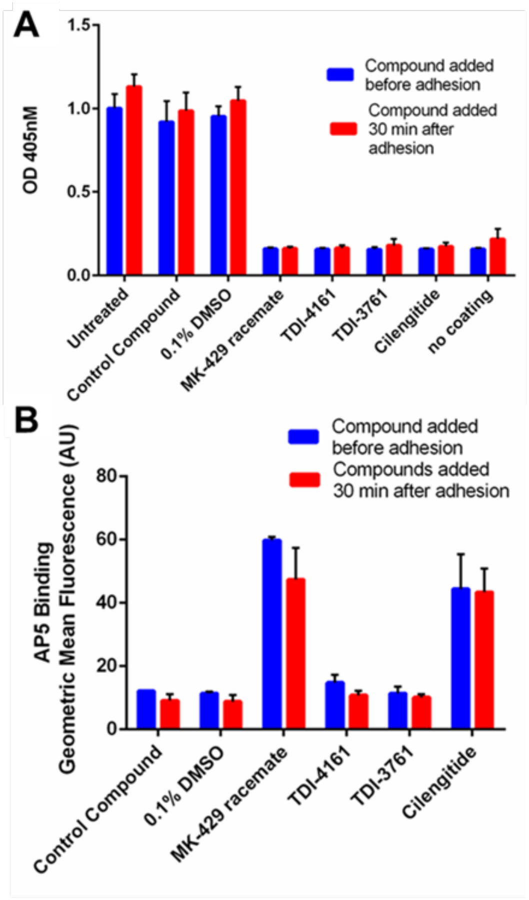
